# Gαs slow conformational transition upon GTP binding and a novel Gαs regulator

**DOI:** 10.1101/2022.10.10.511514

**Authors:** Donghoon Ahn, Davide Provasi, Nguyen Minh Duc, Jun Xu, Leslie Salas-Estrada, Aleksandar Spasic, Min Woo Yun, Juyeong Kang, Dongmin Gim, Jaecheol Lee, Yang Du, Marta Filizola, Ka Young Chung

**Affiliations:** School of Pharmacy, Sungkyunkwan University, Suwon 16419, Republic of Korea; Department of Pharmacological Sciences, Icahn School of Medicine at Mount Sinai, New York, NY 10029, USA; Beijing Advanced Innovation Center for Structural Biology, School of Medicine, Tsinghua University, Beijing 100084, China; Molecular and Cellular Physiology, School of Medicine, Stanford University, Stanford, CA 94305, USA; Department of Biopharmaceutical Convergence, Sungkyunkwan University, Suwon 16419, Republic of Korea; Biomedical Institute for Convergence at SKKU (BICS), Sungkyunkwan University, Suwon 16419, Republic of Korea; School of Life and Health Sciences, Kobilka Institute of Innovative Drug Discovery, Chinese University of Hong Kong, Guangdong 518172, China

**Keywords:** G protein-coupled receptors, G protein, alpha-helical domain, melanoma-associated antigen D2, hydrogen-deuterium exchange mass spectrometry, metadynamics, molecular dynamics, integrative modeling

## Abstract

G proteins are major signaling partners for G protein-coupled receptors (GPCRs). Although stepwise structural changes during GPCR–G protein complex formation and guanosine diphosphate (GDP) release have been reported, no information is available with regard to guanosine triphosphate (GTP) binding. Here, we used a novel Bayesian integrative modeling framework that combines data from hydrogen-deuterium exchange mass spectrometry, tryptophan-induced fluorescence quenching, and metadynamics simulations to derive a kinetic model and atomic-level characterization of stepwise conformational changes incurred by the β_2_-adrenergic receptor (β_2_AR)-Gs complex after GDP release and GTP binding. Our data suggest rapid GTP binding and GTP-induced dissociation of Gαs from β_2_AR and Gβγ, as opposed to a slow closing of the Gαs α-helical domain (AHD). Yeast-two-hybrid screening using Gαs AHD as bait identified melanoma-associated antigen D2 (MAGE D2) as a novel AHD-binding protein, which was also shown to accelerate the GTP-induced closing of the Gαs AHD.

## INTRODUCTION

Heterotrimeric G proteins are a family of guanine nucleotide-binding proteins composed of three distinct subunits (Gα, Gβ, and Gγ). G proteins exist in inactive or active states depending on whether the nucleotide bound to Gα is guanosine diphosphate (GDP) or guanosine triphosphate (GTP). Specifically, GDP-bound Gα forms an inactive trimeric complex with Gβγ, whereas GTP-bound Gα exists in an active state dissociated from both receptor and Gβγ subunits (Milligan and Kostenis, 2006) (Figure 1A).

**Figure 1.**
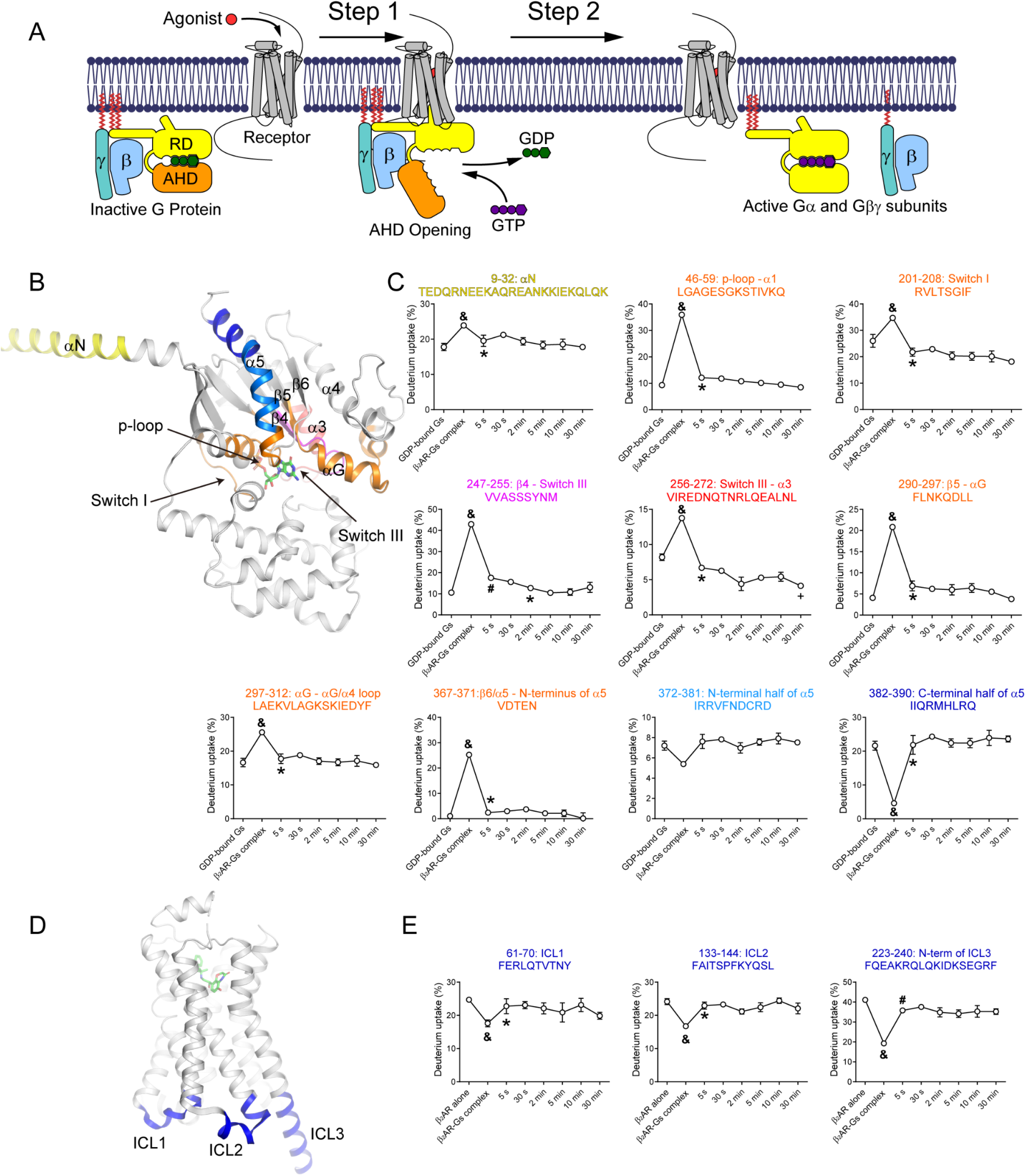
Time-resolved analysis of β_2_AR and Gαs RD after GTPγS addition to the β_2_AR–Gαs complex. (A) G protein activation pathway illustrating the GPCR-mediated GDP release (Step 1), and GTP binding-induced GPCR–G protein complex dissociation and G protein activation (Step 2). RD indicates the Ras-like GTPase domain, and AHD indicates the α-helical domain of the Gα subunit. (B) Regions in the Gαs RD showing HDX profile changes after GTPγS addition color-coded on the X-ray crystal structure of GDP-bound Gs heterotrimer (PDB: 6EG8). Gβγ is omitted for clarity. (C) Pulse-labeling deuterium uptake plots of the peptic peptides color-coded as in (B). (D) Regions in the β_2_AR showing HDX level changes after GTPγS addition highlighted in blue on the X-ray crystal structure of the β_2_AR-Gs complex (PDB: 3SN6). Gs is omitted for clarity. (E) Pulse-labeling deuterium uptake plots of the peptic peptides colored blue as in (D). For (C) and (E), the mass differences >0.3 Da were considered significant. To compare two different time points, a paired t-test was used, and *p* < 0.05 was considered statistically significant. &, the HDX level of the nucleotide-free β_2_AR-Gs complex is significantly different from the HDX levels of GDP-bound Gs or β_2_AR alone. *, the first time point after GTPγS addition when the HDX level returned to that of the GDP-bound Gs or β_2_AR alone. #, the first time point after GTPγS addition when the HDX level was significantly different from that of the nucleotide-free β_2_AR–Gs complex but not yet returned to the levels of GDP-bound Gs or β_2_AR alone. +, first time point showing a statistically significant difference from the time point (*). Error bars represent the standard error of the mean of more than three independent experiments. Data are plotted using a non-linear/non-logarithmic scale. See also **Figures S1 and S2**.

G protein-coupled receptors (GPCRs), one of the largest groups of cell-surface receptors and validated drug targets (Rogers, 2019; Yang et al., 2021), act as guanine nucleotide exchange factors (GEFs) by triggering GDP release from Gα upon their binding to G proteins (Figure 1A). Given the approximately 10-fold higher concentration of GTP than GDP in the cell (Traut, 1994), GTP binding to the empty nucleotide-binding pocket is believed to occur quickly and induce the conformational changes in Gα leading to its dissociation from the receptor and Gβγ and resulting in the full activation of Gα and Gβγ (Figure 1A).

Several high-resolution three-dimensional structures of GPCR–G protein complexes (Draper-Joyce et al., 2018; Garcia-Nafria et al., 2018a; Garcia-Nafria et al., 2018b; Kang et al., 2018; Kato et al., 2019; Koehl et al., 2018; Krishna Kumar et al., 2019; Maeda et al., 2019; Qi et al., 2019b; Rasmussen et al., 2011; Tsai et al., 2019; Tsai et al., 2018; Xing et al., 2020; Zhuang et al., 2020), alongside high-resolution G protein structures bound to GDP or GTP equivalents (Coleman et al., 1994; Liu et al., 2019; Sondek et al., 1994; Sunahara et al., 1997; Wall et al., 1995), have provided an unprecedented level of atomistic detail into GPCR–G protein signaling (see examples in Figure S1A–C). These structures have revealed that the nucleotide-binding pocket is located between the Ras-like GTPase domain (RD) and the α-helical domain (AHD) of Gα (Preininger et al., 2013; Wall *et al*., 1995) (Figure 1A and Figure S1A). GPCRs bind to the Gα RD, particularly at the α5 helix and a hydrophobic core formed by the αN–β1 hinge, β2–β3, and α5, mostly through interactions with their cytosolic core and intracellular loops (ICLs) (Figure S1B) (Garcia-Nafria and Tate, 2019; Glukhova et al., 2018). Notably, these interactions have been suggested to trigger allosteric conformational changes near the nucleotide-binding pocket, which lead to GDP release and the opening of the Gα AHD with respect to the RD (Garcia-Nafria and Tate, 2019; Glukhova *et al*., 2018; Preininger *et al*., 2013) (Figure 1A and Figure S1B). In contrast to the large differences between the GDP-bound and nucleotide-free receptor-bound Gα states (compare Figure S1A with Figure S1B), GDP-bound and GTP-bound Gα showed very little structural discrepancies, which are mostly limited to Switch II rearrangements at the Gβγ-binding interface (Figure S1C).

Notwithstanding the level of molecular detail revealed by these structures, the complete series of time-resolved conformational transitions undergone by Gα during its activation process is unclear. Given the importance of these transitions, they have been the focus of extensive research in recent years, using a variety of biophysical, biochemical, and computational approaches (Ahn et al., 2021; Dror et al., 2015; Du et al., 2019; Ham et al., 2021; Kato *et al*., 2019; Kim et al., 2021; Liu *et al*., 2019; Van Eps et al., 2011; Westfield et al., 2011). Notably, both computational (Sun et al., 2018) and experimental (Du *et al*., 2019) studies support the existence of alternative transient intermediates to the reported high-resolution structures during receptor-induced GDP release and GPCR–G protein complex formation (Figure 1A, step 1). In particular, our previous experimental study using hydrogen/deuterium exchange mass spectrometry (HDX-MS) in a time-resolved manner (pulse-labeling HDX-MS) revealed the stepwise conformational changes during β_2_-adrenergic receptor (β_2_AR)–Gs complex formation and GDP release, which confirmed that the early β_2_AR–Gs complex adopts a transient conformation that is different from the reported X-ray crystal structure of the nucleotide-free β_2_AR–Gs complex (PDB: 3SN6) (Du *et al*., 2019).

Here, we used pulse-labeling HDX-MS (Figure S1D) to investigate the time-resolved conformational changes incurred by β_2_AR and Gs after guanosine 5’-O-[γ-thio]triphosphate (GTPγS), a non-hydrolyzable GTP analog, was added to the nucleotide-free β_2_AR–Gs complex (Figure 1A, step 2). We further analyzed the movement of the Gαs AHD by monitoring its separation from the Gαs RD using a tryptophan-induced fluorescence quenching (TrIQ) technique. These data were then integrated with the results of well-tempered metadynamics simulations (Barducci et al., 2008) using a novel integrative modeling framework that combines notions of Bayesian inference (Rout and Sali, 2019) with the Maximum Caliber (MaxCal) principle (Dixit and Dill, 2014; Dixit et al., 2015), Markov State Modeling (MSM) (Perez-Hernandez et al., 2013; Schwantes and Pande, 2013), and transition path theory (TPT) (E and Vanden-Eijnden, 2010). The goal was to translate experimental observations of conformational changes incurred by the β_2_AR–Gs complex after GTPγS addition into structural ensembles of the most probable metastable states sampled by Gαs upon GTPγS binding and their kinetic relationships. Finally, we used yeast-two-hybrid (Y2H) library screening to identify Gαs AHD-binding proteins that regulate the kinetics during the GTP binding-induced closing of the Gαs AHD.

## RESULTS

### Rapid GTPγS binding to Gαs

HDX-MS translates measurements of the exchange between amide hydrogens in the protein backbone and deuterium in the solvent in terms of the stability of the protein’s secondary structure as well as its solvent accessibility (Harrison and Engen, 2016). Accordingly, HDX-MS can provide conformational information about the receptor–G protein interfaces, nucleotide-binding pocket, and GTP-induced allosteric conformational changes, albeit not at the single-residue or atomic-level resolution. To analyze the stepwise conformational changes incurred by the β_2_AR–Gs complex upon GTPγS binding, we prepared the nucleotide-free β_2_AR–Gs complex for pulse-labeling HDX-MS experiments, as described in the STAR Methods (Figure S1D). Briefly, we collected aliquots of the agonist-bound β_2_AR, GDP-bound Gs, and nucleotide-free β_2_AR–Gs complex, as well as aliquots of protein samples at specific time points (5 s, 30 s, 2 min, 5 min, 10 min, and 30 min) after GTPγS addition to the nucleotide-free β_2_AR-Gs complex. The collected samples were immediately exposed to a D_2_O pulse for 10 s (Figure S1D). The detailed HDX-MS analysis parameters and mass spectrum information of all analyzed peptides are summarized in Supplementary Data.

The HDX levels were higher for peptides near the nucleotide-binding pocket of Gαs (p-loop through α1, Switch I, Switch III, β5 through the αG/α4 loop, and the β6/α5 loop through the N-terminus of α5) in the nucleotide-free β_2_AR–Gs complex compared with those of GDP-bound Gs (Figure 1B, magenta-, red-, and orange-colored regions; Figure 1C, compare the first two time points in plots of peptides 46–59, 201–208, 247–255, 256–272, 290–297, 297–312, and 367–371), reflecting increased solvent accessibility and structural dynamics after GDP release, which is consistent with our previous reports (Chung et al., 2011; Du *et al*., 2019; Kim et al., 2020).

Within 5 s of adding GTPγS to the nucleotide-free β_2_AR–Gs complex, the HDX levels of all these peptides, except for the Switch III peptide 247–255, returned to the HDX levels of the GDP-bound Gs, while the HDX levels of the Switch III peptide 247– 255 returned to approximately 80–90% of the HDX levels of GDP-bound Gs (Figure 1C), suggesting rapid binding of GTPγS at the nucleotide-binding pocket. While the HDX levels of peptides 46–59, 201–208, 290–297, 297–312, and 367–371 of Gαs did not change significantly after 5 s (Figure 1C; orange-colored regions in Figure 1B), the HDX levels of the Switch III peptide 256–272 continued to change during 30 min, becoming significantly lower than those of GDP-bound Gs at the end of the experiment (Figure 1C; red-colored region in Figure 1B). Similarly, the HDX levels of Switch III peptide 247–255 continued to decrease after their drastic change within 5 s, but for only 2 min, ultimately returning to the levels of the GDP-bound Gs (Figure 1C; magenta-colored region in Figure 1B). In summary, the pulse-labeling HDX-MS data collected for peptides near the nucleotide-binding pocket suggested rapid GTPγS binding (within 5 s) and additional prolonged conformational changes at Switch III after GTPγS binding.

### Rapid dissociation of Gαs from β_2_AR and Gβγ after GTPγS binding

The HDX levels were lower at the β_2_AR–Gs interface (C-terminal half of α5 of Gαs and ICLs 1, 2, and 3 of β_2_AR) in the nucleotide-free β_2_AR–Gs complex compared to the GDP-bound Gs (Figure 1B, blue-colored regions; Figure 1C, compare the first two time points in the plot of peptide 382–390), as well as compared with the levels of the agonist-bound β_2_AR alone (Figure 1D, blue-colored regions; Figure 1E, compare the first two time points in all plots). This suggested reduced solvent accessibility and stabilization of the secondary structures at the β_2_AR–Gs interface, which is consistent with our previous reports (Chung *et al*., 2011; Du *et al*., 2019; Kim *et al*., 2020). The HDX levels of the peptide from the N-terminal half of α5 also showed a similar trend, although it was not significant (less than 0.3 Da differences) (Figure 1B, light blue-colored region; Figure 1C, peptide 372–381). The HDX levels of αN were higher in the β_2_AR–Gs complex compared with those detected in the GDP-bound Gs (Figure 1B, yellow-colored region; Figure 1C, compare the first two time points in the plot of peptide 9–32), suggesting that αN undergoes conformational changes upon β_2_AR–Gs complex formation. Within 5 s of GTPγS addition, the HDX levels of these regions returned to those of GDP-bound Gs (Figures 1C, peptides 9–32 and 382–390) or the agonist-bound β_2_AR alone (Figure 1E), suggesting that GTPγS-bound Gαs dissociates rapidly from the receptor.

Pulse-labeling HDX-MS analysis of Gβγ revealed rapid conformational changes in Gβ WD1 S1, Gβ WD1 S3 through S4, and Gβ WD2 S2 after GTPγS addition (Figure S2). The HDX levels of Gβ WD1 S1 were higher in the nucleotide-free β_2_AR–Gs complex than those of GDP-bound Gs and quickly (within 5 s) returned to the level of the GDP-bound Gs after GTPγS addition (Figure S2, orange-colored region, peptide 33–46). The HDX levels of Gβ WD1 S3 through S4 and those of WD2 S2 did not differ between GDP-bound Gs and the nucleotide-free β_2_AR–Gs complex (Figure S2, cyan-colored regions, compare the first two time points in plots of peptides 81–99 and 111– 118); however, upon GTPγS addition to the β_2_AR–Gs complex, these regions showed increased HDX levels within only 5 s, which plateaued after 30 s (Figure S2, cyan-colored regions, peptides 81–99 and 111–118). These HDX level changes in peptides 81–99 and 111–118 of Gβγ likely reflect conformational changes due to the dissociation of Gβγ from Gαs, given that they were observed only after GTPγS addition but not upon β_2_AR–Gs complex formation. Overall, these results suggest that Gβγ may also quickly dissociate from GTPγS-bound Gαs.

### Slow conformational changes of the Gαs AHD after GTPγS binding

A notable observation of the pulse-labeling HDX-MS analysis was that several Gαs AHD peptides showed slow changes in the HDX levels upon GTPγS binding. The HDX levels of these peptides were higher in the nucleotide-free β_2_AR–Gs complex compared with those in GDP-bound Gs (Figure 2A, compare the first two time points in all plots), reflecting the AHD displacement and increased local conformational dynamics in the nucleotide-free β_2_AR–Gs complex compared to the GDP-bound Gs (compare Figure S1A and Figure S1B).

**Figure 2.**
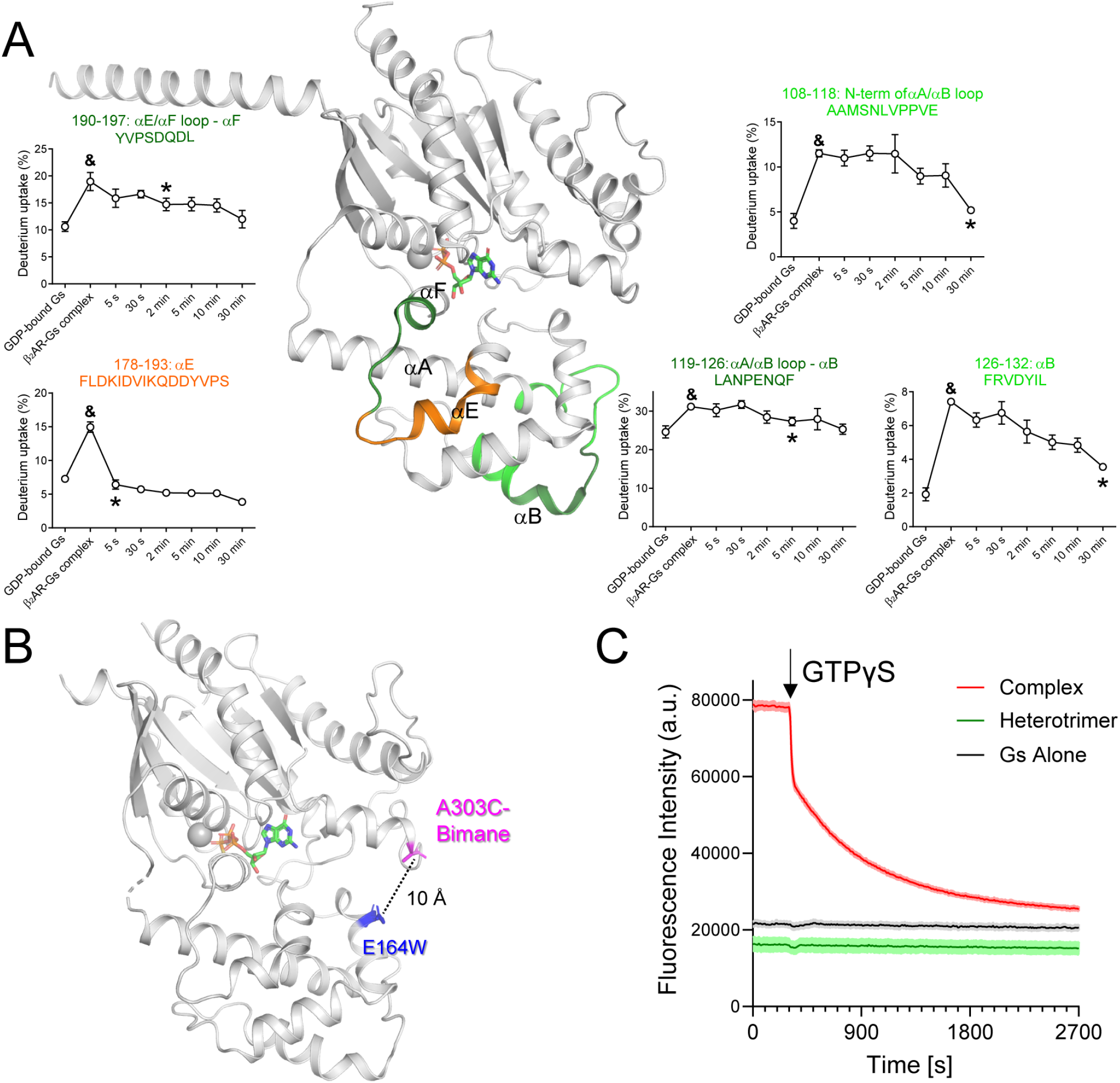
Time-resolved analysis of conformational changes of the Gαs AHD after GTPγS addition to the β_2_AR–Gαs complex. (A) Regions in the Gαs AHD showing HDX level changes after GTPγS addition color-coded on the X-ray crystal structure of the GDP-bound Gs heterotrimer (PDB: 6EG8). Gβγ is omitted for clarity. Pulse-labeling deuterium uptake plots are shown for the color-coded peptic peptides. The mass differences >0.3 Da were considered significant. To compare two different time points, a paired *t*-test was used, and *p* < 0.05 was considered statistically significant. &, the HDX level of the nucleotide-free β_2_AR-Gs complex is significantly different from that of GDP-bound Gs. *, the first time point after GTPγS addition when the HDX level returned to that of the GDP-bound Gs. Error bars represent the standard error of the mean of more than three independent experiments. Data are plotted using a non-linear/non-logarithmic scale. (B) Residues labeled with bimane (magenta) or mutated to Trp (blue) indicated as sticks on the X-ray crystal structure of GTPγS-bound Gαs (PDB: 1AZT). (C) Time-resolved tryptophan-induced bimane quenching analysis of the β_2_AR–Gs complex after GTPγS addition. The data are representative of three independent experiments. See also **Figures S1 and S3**.

Except for the αE region, whose HDX levels quickly returned to those of the GDP-bound Gs within 5 s of GTPγS addition (Figure 2A, orange-colored region, peptide 178–193), the HDX levels of other peptides depicted in Figure 2A showed slow and continued changes (peptides 108–118, 119–126, 126–132, and 190–197). The N-terminus of the αA/αB loop and αB slowly returned to the HDX levels of GDP-bound Gs within 30 min (Figure 2A, green-colored region, peptides 108–118 and 126–132). The αA/αB loop through αB and the αE/αF loop through αF returned to the HDX levels of GDP-bound Gs after 2–5 min (Figure 2A, dark green-colored regions, peptides 119–126 and 190–197). These results suggested that the Gαs AHD undergoes relatively slow and continued conformational changes even after GTPγS binding to Gαs and its consequent dissociation from the receptor and Gβγ (within 5 s in the current experimental system, as shown in Figure 1B–E).

### Slow AHD closing kinetics after GTPγS binding

To understand whether the slow HDX changes in the Gαs AHD (Figure 2A) are related to the slow AHD closing kinetics, we developed an experimental system that monitors the separation between the RD and AHD using TrIQ (Figure 2B). The bimane fluorescence is quenched when aromatic residues such as Trp, Tyr, and Phe are nearby (Jones Brunette and Farrens, 2014). Therefore, given their vicinity (α-carbon atoms within 10 Å) in the GTPγS-bound Gαs crystal structure (PDB: 1AZT), we engineered Gαs to replace residue A303 in the RD with Cys for monobromobimane labeling and residue E164 in the AHD with Trp (Figure 2B). In this construct, the bimane fluorescence is expected to be quenched in the nucleotide-bound closed Gαs conformation (see Figure S1A and S1C) but unquenched in the nucleotide-free open conformation (see Figure S1B). As there are other Cys residues in Gαs, we mutated all solvent-exposed Cys residues (Figure S3A, red-colored residues) to Ser to prevent unwanted bimane labeling, and confirmed that these mutations do not affect the function of Gαs by measuring successful BODIPY-FL-GTPγS uptake profiles (Figure S3B) and heterotrimer formation with Gβγ (Figure S3C).

We first verified the suitability of this construct for monitoring the separation between the RD and AHD by recording (a) similar bimane fluorescence values between GDP-bound and GTPγS-bound Gαs (Figure S3D), in line with their conformational states being closed (Figure S1C); and (b) higher bimane fluorescence values for the nucleotide-free β2AR–Gs complex compared to the GDP-bound Gs heterotrimer or GDP-bound Gαs (Figure S3E), in line with AHD being displaced in the nucleotide-free β_2_AR–Gs complex (Figure S1B).

We then used this TrIQ method to monitor Gαs AHD closing in a time-resolved manner. After GTPγS addition to the nucleotide-free β_2_AR–Gs complex, the bimane fluorescence on residue A303C decreased in two steps: an initial fast decrease (with a time constant of 50 ± 8 s), followed by a slow (time constant of 655 ±10 s) and continued decrease (Figure 2C). This observation suggested that the AHD may adopt long-lived intermediate conformational states during its GTPγS binding-induced closing.

### Simulated conformational states of Gαs after GDP release and GTP binding

To obtain an atomic-level characterization of Gαs conformations after GDP release and GTP binding (Figure 1A, step 2), we simulated Gαs either starting from its closed, GTPγS-bound crystal structure (PDB: 1AZT (Sunahara *et al*., 1997)) in the presence of a bound Mg^2+^ ion and a water environment or starting from its nucleotide-free conformation in the β2AR–Gs complex (PDB: 3SN6 (Rasmussen *et al*., 2011)) in a hydrated 1-palmitoyl-2-oleoyl-sn-glycero-3-phosphocholine (POPC)/10% cholesterol lipid bilayer (see the STAR Methods for details of the simulated systems). Since standard molecular dynamics (MD) simulations would require very long simulation times to reproduce the large conformational changes between the closed and open states of Gαs, after equilibration of the GTPγS–Gαs and β_2_AR–Gs systems, we used well-tempered metadynamics (Barducci *et al*., 2008) to enhance the sampling of the separation and relative orientation between their RD and AHD regions using the two collective variables (CVs) illustrated in Figure S4A (see the STAR Methods for simulation details).

The reconstructed free-energy surfaces from simulations are illustrated in Figure S4B. Both the GTPγS–Gαs and nucleotide-free β2AR–Gs systems explored a wide range of relative orientations of the RD and AHD, with different degrees of separation between the two domains. The GTPγS–Gαs system explored two main conformational ensembles: one encompassing closed AHD states (CV1 ~30°, CV2 ≲ 2) and one characterized by open AHD states with large values of the contact map distance (CV2 ≳ 10), as well as a wide range of angles (60°≲ CV1 ≲ 200°). The two ensembles exhibited similar free energy but were separated by a free-energy barrier above 10 kT (Figure S4B). Notably, despite the separation between the RD and AHD, and in agreement with previous computational studies (Dror *et al*., 2015; Sun *et al*., 2018), GTPγS remained firmly bound to the protein throughout the simulation, suggesting that the Gαs AHD can exist in the open state even when GTPγS is bound to Gαs.

In simulations of the nucleotide-free β_2_AR–Gs system, the presence of the receptor and Gβγ appeared to stabilize an ensemble of Gαs states with similar degrees of RD–AHD separation to the one observed in the GTPγS–Gαs system. However, it showed more heterogeneous distributions of interdomain contacts compared with those of the GTPγS-bound Gαs crystallographic closed state (PDB: 1AZT) (CV1 ~30°; 2 ≲ CV2 ≲ 10) or the open state sampled by the GTPγS-bound Gαs system (60° ≲ CV1 ≲ 200°; 7 ≲ CV2 ≲ 15). In the absence of GTPγS, the ensemble of Gαs open states (CV1 > 60°) was energetically preferred over that of the Gαs closed states, with a free-energy difference of 1.28 kT in the case of the nucleotide-free β_2_AR–Gs system and ~0 kT in the GTPγS-bound Gαs system (Figure S4B).

### Atomic-level kinetic model of Gαs activation

Although the metadynamics simulations offered insight into the conformational states that are preferably adopted by Gαs in its nucleotide-free form within the β_2_AR–Gs complex or GTPγS–Gαs, the long timescales needed for nucleotide binding and unbinding, as well as the bias introduced by the metadynamics simulations, limit the type of information that can be directly extracted by such analyses. To obtain an atomic-level description of Gαs activation steps after GDP release and GTP binding (Figure 1A, step 2) consistent with the time-resolved conformational changes of the Gαs backbone at the second-to-minute timescale from the pulse-labeling HDX-MS (see STAR Methods and Figures 1C and 2A), as well as the Gαs conformational states inferred by TrIQ at time zero and infinite (equilibrium) (Figure 2C), we integrated these data with predictions of average residue protection factors and fluorescence quenching computed for each conformational cluster derived from the well-tempered metadynamics simulations (Barducci *et al*., 2008) of the GTPγS-bound Gαs and nucleotide-free β_2_AR– Gs systems.

Specifically, we developed a novel integrative modeling framework that combines the notions of Bayesian inference (Rout and Sali, 2019), the MaxCal principle (Dixit and Dill, 2014; Dixit *et al*., 2015), MSM (Perez-Hernandez *et al*., 2013; Schwantes and Pande, 2013), and TPT (E and Vanden-Eijnden, 2010). Accordingly, the experimental and computational data were integrated by deriving a Markov transition matrix of the probabilities of moving from one metadynamics-derived conformational cluster to another. Specifically, the posterior probability of the proposed kinetic model *M* was expressed as the posterior distribution *p*(*M*|*O*) ∝ *p*(*O*|*M*)*p*(*M*), where *p*(*O*|*M*) is the likelihood that model *M* would predict the observed experimental data *O* and *p*(*M*) is the prior probability of the model before observing *O*, in which the probability of the transition matrix is assigned by MaxCal (Dixit and Dill, 2014; Dixit *et al*., 2015) while priors of the equilibrium probabilities correspond to the metadynamics-derived free energies (see STAR Methods for details). TPT analysis of the transition matrix further enriched this scheme by providing estimates for the mean first passage times of the transitions between the ensembles of experimentally weighted conformational populations of Gαs bound to GTPγS or within the nucleotide-free β_2_AR–Gs complex derived from the metadynamics simulations.

In addition to the TrIQ data displayed in Figure 2C, the deuterium uptake fractions of the 15 protein regions spanning the RD and AHD of Gαs illustrated in Figures 1C and 2A were used as the observable data *O* in the aforementioned integrative modeling approach. An established approach that enables estimating HDX protection factors for a given protein conformation (Best and Vendruscolo, 2006) was used to link the HDX-MS experimental data to the conformations of the GTPγS–Gαs and nucleotide-free β_2_AR–Gs systems sampled by metadynamics simulations. After k-means clustering of the metadynamics trajectories using pairwise distances between every third Cα atom in Gαs projected onto the first 10 tICA components, a set of 16 conformational macrostates, *R_s_* (see STAR Methods for details), was used to compute average residue protection factor estimates for each of the 15 protein regions illustrated in Figures 1C and 2A using 10 frames of each conformational cluster (see STAR Methods). Similarly, these cluster conformations were used to predict fluorescence quenching using a simple contact model (Jones Brunette and Farrens, 2014).

The proposed Bayesian integrative modeling framework indicated that very good agreement could be achieved between predicted and measured values of the deuterium uptake and fluorescence quenching for 13 out of 15 regions (Figure S5), with peptides 9–32 and 247–255 showing the worst agreement. Only values of the 13 peptides with the best agreement between predicted and experimental measurements were used as observable data for the kinetic model described below.

Figure 3A illustrates the proposed kinetic model of Gαs activation upon GTPγS binding, with nodes of the network representing highly populated states (i.e., states whose maximum probability over time is larger than 10%; see values in Table S1) at either time zero or equilibrium (gray and black circles, respectively, in Figure 3A), and the connections among them indicating the timescales needed to transition from one state to another. Initially (t = 0), Gαs mostly explored two nucleotide-free conformational states when bound to β_2_AR and Gβγ, herein labeled states 10 and 13 with a probability of 29% and 54%, respectively (Table S1), which are characterized by different degrees of separation between the RD and AHD (mean distance of 65.1 Å or 49.4 Å between A303 and E164, respectively) (Figure 3A). This is consistent with the observation that the AHD in the nucleotide-free β_2_AR–Gs complex is mobile rather than assuming a fixed position (Westfield *et al*., 2011).

**Figure 3.**
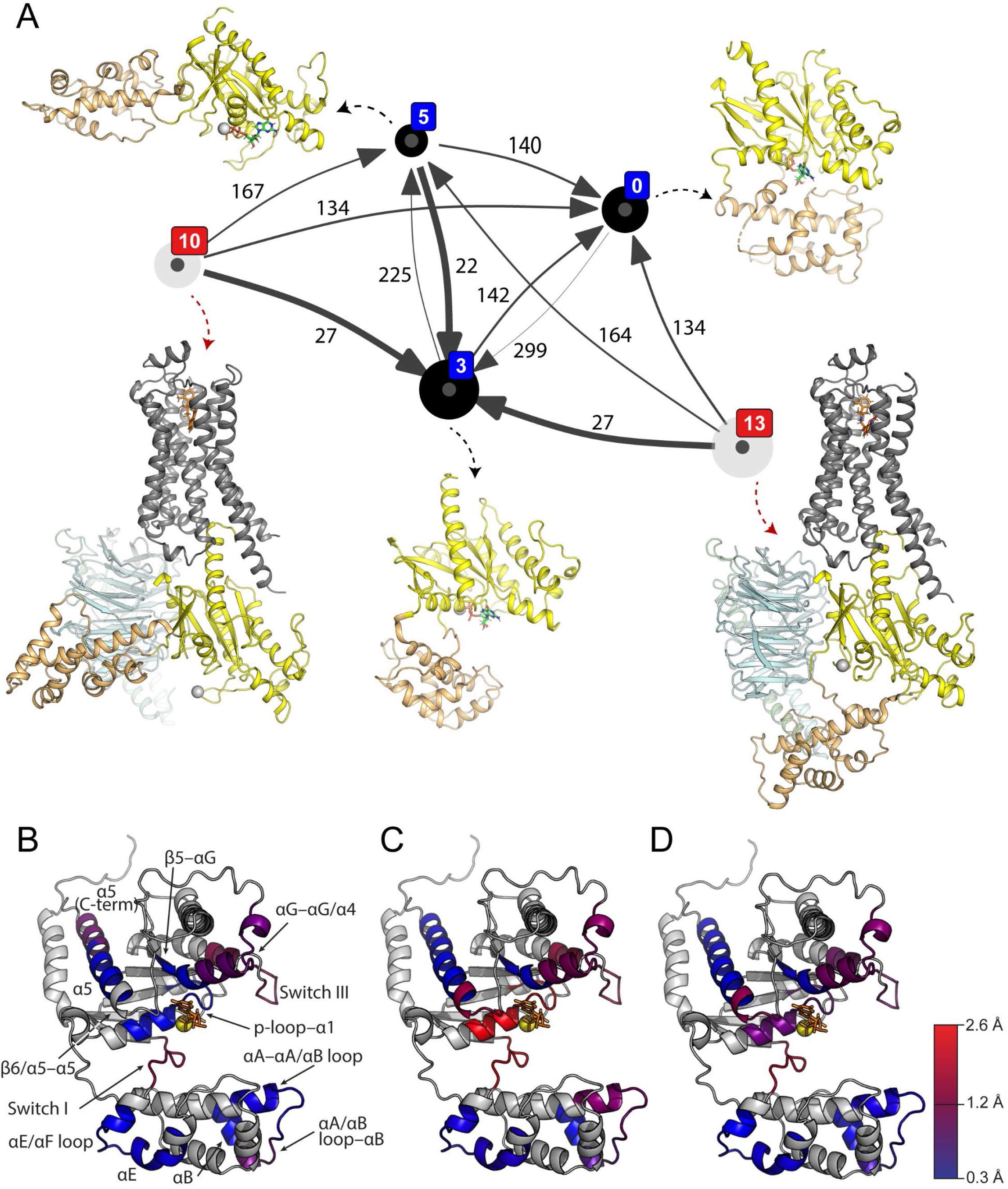
Highly probable conformational states explored by Gαs after GTPγS addition according to the proposed kinetic model. (A) Highly populated states (maximum probability > 10%) in the GTPγS–Gαs simulation are indicated by blue numbers, while states in the nucleotide-free β_2_AR–Gs simulation are indicated by red numbers. The size of black and gray circles is proportional to the probability of a certain state to be populated at equilibrium (long time scales) or time zero, respectively. Arrows represent mean transition timescales between states, with thick and thin arrows indicating fast and slow transition times, respectively. Mean first passage times (MFPT, in seconds) are reported on the arrows, and their credible intervals are reported in Table S2. Representative structures of each highly populated state are shown in cartoon representations with the receptor in gray, RD of Gα in yellow, AHD of Gα in light orange, Gβ in light blue, and Gγ in light green. Magnesium ion (Mg^2+^) is indicated with a gray sphere, and, where present, GTPγS is shown as sticks. (B–D) Comparison of the RMSD values of the HDX-MS peptic peptides between macrostates 3 and 0 (B), 3 and 13 (C), and 3 and 10 (D), respectively, color-coded from low (blue) to high (red) values on a representative structure of state 3. See also **Figures S4, S5, and S8, as well as Tables S1–S3**.

In the presence of GTPγS, protein states 10 and 13 most rapidly (mean first passage times (MFPTs) of ~30 s; see Table S2) transitioned to a highly populated (equilibrium probability of 51%) GTPγS-bound conformation with an open AHD (state 3, with an inter-domain mean A303–E164 distance of 28.7 Å) (Figure 3A, thick arrows starting from state 10 or 13 to state 3). Slower transitions were recorded from states 10 and 13 to a lower-probability open GTPγS-bound conformation state 5 (equilibrium probability of 13% and inter-domain mean A303–E164 distance of 77.3 Å; reached, on average, in ~160 s) or to a closed conformational state 0 (equilibrium probability of 29% and an inter-domain mean A303–E164 distance of 9.8 Å; reached, on average, in ~130 s) (Figure 3A, thin arrows starting from state 10 or 13 to state 5 or 0; and Tables S1– S2). State 0 closely resembled the X-ray crystal structure of GTPγS-bound Gαs that was used as a starting point for simulations (RMSD of 1.22 Å from PDB: 1AZT).

The half-life of state 3 between the nucleotide-free Gαs conformation bound to both β_2_AR and Gβγ and the closed GTPγS-bound conformational state of Gαs was estimated to be 142 s, which is shorter than the half-life of the closed GTPγS-bound conformational state (state 0; 299 s), but much longer than the half-life of the alternative open GTPγS-bound state (state 5; 22 s) (Figure 3A). Notably, state 5 rapidly converted into state 3 (22 s), but it took much longer for the AHD to transition from state 3 to state 5 (225 s) (Figure 3A). The mean distance between RD A303 and AHD E164 in state 3 (28.7 Å) was smaller than that found in states 10, 13, or 5 (65.1 Å, 49.4 Å, and 77.2 Å, respectively) but larger than that of state 0 (9.8 Å). These results suggest that state 3 may represent the long-lived intermediate state inferred by the data shown in Figure 2C.

Notably, state 3 differed from the X-ray crystal structure of GTPγS-bound Gαs, as well as the two most populated states (10 and 13) of the nucleotide-free Gαs conformation bound to the β2AR and Gβγ, not only in terms of mean distance separation between RD A303 and AHD E164 but also for the relative orientation between these two domains. RMSD comparison between the identified highly populated long-lived intermediate state 3 of GTPγS-bound Gαs with the identified closest states to the GTPγS-bound crystal structure 1AZT (state 0) or with the highly probable nucleotide-free Gαs conformations bound to the β_2_AR and Gβγ (states 10 and 13) revealed specific regions that are, on average, more different than others (Table S3 and Figure 3B–D). However, in general, the differences were larger between states 3 and 10 or 13 (Figure 3C–D) than between states 3 and 0 (Figure 3B). Specifically, regions 46– 59 (p-loop–α1), 201–208 (Switch I), and 256–272 (Switch III) exhibited the largest differences between states 3 and 13 (Figure 3C), with Switch I also differing the most between states 3 and 10 (Figure 3D). Although the largest differences (RMSD > 2.0) were found between states 3 and 13, pertaining to p-loop–α1, Switch I, and Switch III, the latter two also deviated significantly (RMSD > 1.5) in state 3 compared to states 10 and 0.

### Novel Gαs AHD-binding protein facilitating AHD closing

The G-protein signaling cycle is regulated by several proteins, including GEFs (e.g., GPCRs, GIV/Girdin, or Ric-8A) and GTPase-activating proteins [GAPs; e.g., regulators of G protein signaling (RGS)] (Kach et al., 2012; Kalogriopoulos et al., 2019; Siderovski and Willard, 2005; Srivastava and Artemyev, 2020; Srivastava et al., 2019). As our data support slow AHD closing enabled by the long-lived intermediate state of Gαs after GTPγS binding, we searched for an AHD-binding protein that might regulate the AHD closing kinetics. To this end, we employed Y2H library screening with the Gαs AHD as bait and identified melanoma-associated antigen D2 (MAGE D2, NM_014599) as a novel Gαs AHD-binding protein (Figure S6A).

Mutations in MAGE D2 have been reported to cause Bartter’s syndrome, a rare autosomal recessive renal tubular disorder (Laghmani et al., 2016). In the same study, MAGE D2 was discovered to interact directly with Gαs, and other studies suggested that MAGE D2 modulates GPCR signaling (Laghmani *et al*., 2016; Paek et al., 2017; Reusch et al., 2022).

In this study, we identified that AHD of Gαs is the binding site of MAGE D2. The overexpressed FLAG-tagged Gαs AHD was co-localized with endogenously expressing MAGE D2 in HEK293T cells (Figure 4A). Moreover, turboGFP-tagged MAGE D2 were co-immunoprecipitated with FLAG-tagged Gαs AHD in HEK293T cells (Figure 4B). These results suggest that AHD of Gαs can interact with MAGE D2 in the cellular context.

**Figure 4.**
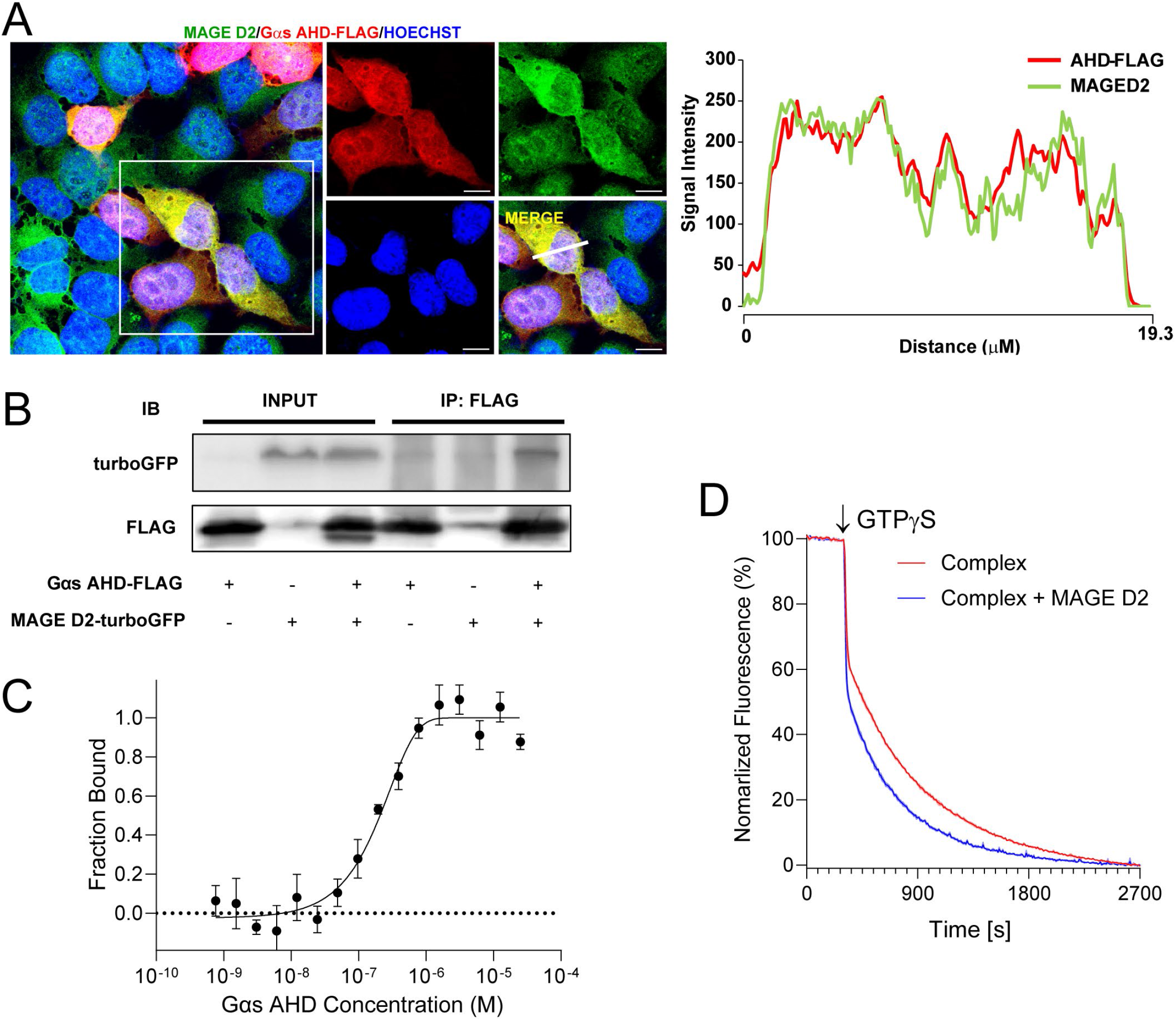
Novel Gαs AHD-binding protein that facilitates AHD closing. (A) Representative confocal images of HEK293T cells transiently expressing Gαs AHD-FLAG. HEK293T cells were stained with antibodies against FLAG (red) and MAGE D2 (green). Blue represents Hoechst 33342. Right: Representative co-localization tracer profile along the line indicated in the inset. Scale bar, 10 μm. (B) Immunoblot (IB) analysis of FLAG immunoprecipitates (IP line) and cell lysates (Input line) from HEK293T cells transiently co-expressing Gαs AHD-FLAG and MAGE D2-turboGFP. (C) Binding curve of the Gαs AHD with the MAGE D2 MHD analyzed by microscale thermophoresis (MST). A titration series of the Gαs AHD was performed, while the concentration of fluorescence-labeled MAGE D2 MHD was fixed (see STAR Methods for details). Error bars represent the standard error of the mean of more than three independent experiments. (D) Time-resolved tryptophan-induced bimane quenching analysis of the β_2_AR–Gs complex with or without the MAGE D2 MHD after GTPγS addition. The data are representative of three independent experiments. See also **Figures S3, S6, and S7**.

The direct interaction between MAGE D2 and Gαs AHD was confirmed using purified proteins and microscale thermophoresis (MST) analysis (Figure 4C). Since all MAGE proteins have a conserved domain called the MAGE homology domain (MHD) that consists of two winged-helix motifs (WH-A and WH-B), which is reported as a protein–protein interaction site (Doyle et al., 2010; Newman et al., 2016), we generated a truncated MAGE D2 construct that only contains the MHD to facilitate protein purification (Figure S6B). MST analysis showed that the purified Gαs AHD interacts with purified MAGE D2 MHD with a K_D_ of 155 ± 47 nM.

We then tested if MAGE D2 binding affects the AHD closing kinetics using the TrIQ technique illustrated in Figure 2B. GTPγS was added to the nucleotide-free β2AR–Gs complex with or without co-incubation of MAGE D2 MHD, and the bimane fluorescence at residue A303C was monitored. Co-incubation of MAGE D2 MHD accelerated the decrease of bimane fluorescence, suggesting that binding MAGE D2 accelerates the closing of the Gαs AHD (Figure 4D).

## DISCUSSION

This study describes the sequential conformational transitions incurred by a GPCR–G protein complex upon GTPγS binding, using the β_2_AR–Gs complex as a model system. Importantly, we provide the first atomic-level structural interpretation of the stepwise conformational changes the β_2_AR–Gs complex experiences after GDP release and GTPγS binding, using a novel Bayesian integrative modeling framework that combines experimental observations of GTPγS-induced β_2_AR–Gs complex dissociation and closing of the Gαs AHD from HDX-MS and TrIQ data with structural ensembles of β_2_AR–Gs and Gαs sampled by metadynamics. Taken together, our data suggest (i) rapid GTPγS binding to the Gαs nucleotide-binding pocket (Figure 5, step i), (ii) rapid Gαs dissociation from the receptor and Gβγ (Figure 5, step ii), and (iii) slow closing of the Gαs AHD and the existence of a long-lived intermediate state (Figure 5, step iii-a).

**Figure 5.**
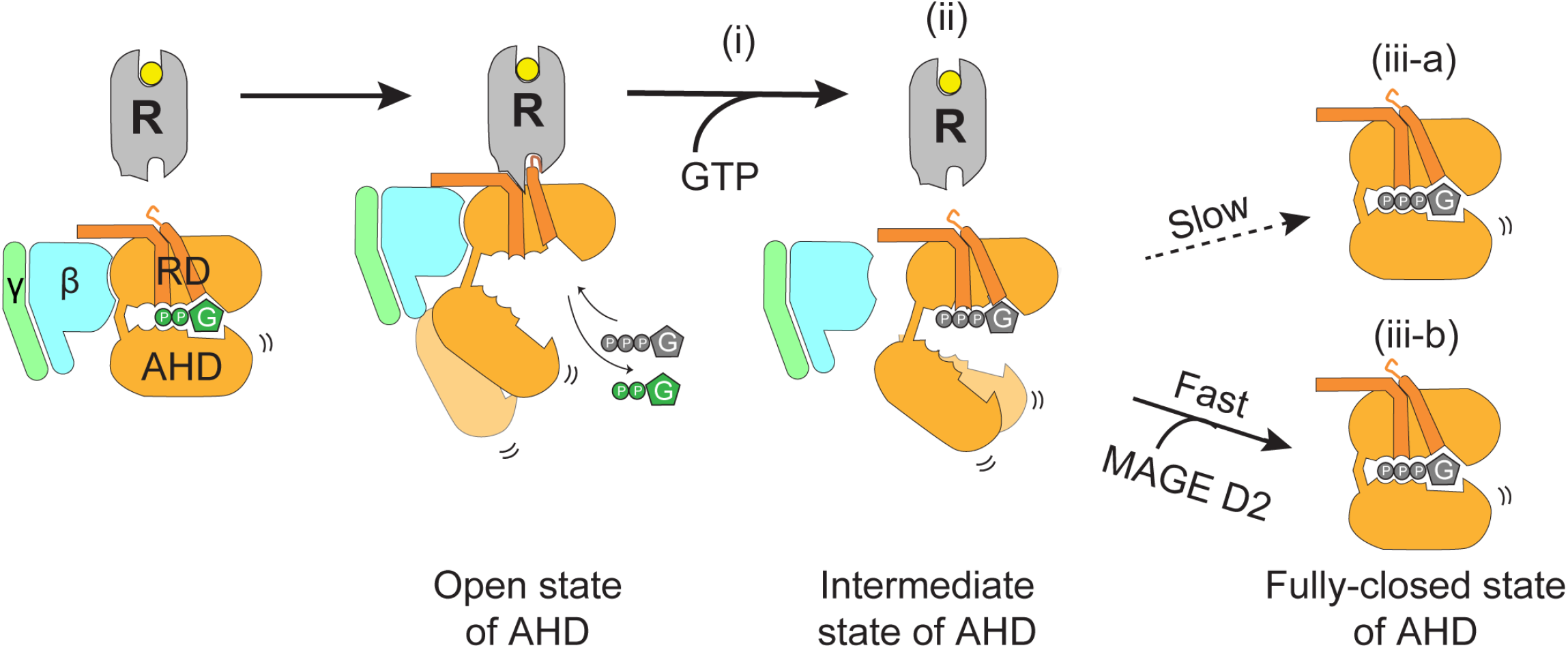
Summary cartoon illustrating the proposed sequence of events after GTP binding to a GPCR–Gs complex with or without MAGE D2. An agonist-activated receptor (R) induces GDP release from Gs, and GTP quickly binds to the empty nucleotide-binding pocket (step i). Upon binding of GTP, Gαs rapidly dissociates from the receptor and Gβγ (step ii). The AHD adopts a long-lived intermediate state (step ii) until it is fully closed, a process that occurs slowly (step iii-a). Closing of the AHD is accelerated in the presence of MAGE D2 (step iii-b). RD indicates the Ras-like GTPase domain, and AHD indicates the α-helical domain.

Our proposed kinetic model identified highly populated conformational ensembles of both the GTPγS-bound Gαs and the nucleotide-free Gαs in complex with β_2_AR and Gβγ, with alternative orientations and separations of the AHD with respect to the RD compared with those observed in available high-resolution crystal structures. Moreover, the model provided estimates of the transition time scales between these states. On the one hand, the observed AHD conformational variability was expected in light of previous reports stating that the AHD could either not be resolved at high resolution or assumed different orientations in X-ray crystallography or cryo-electron microscopy structures of GPCR–G protein complexes (Duan et al., 2021; Rasmussen *et al*., 2011; Sun et al., 2021; Zhang et al., 2020), as well as inferences from double electron-electron resonance spectroscopy (Van Eps *et al*., 2011) and molecular dynamics simulations (Dror *et al*., 2015). On the other hand, the kinetic model of the conformational transitions of Gαs upon GTPγS binding and the atomic details of the most populated conformational states sampled by Gαs after GTP binding were unknown until now. Among the most populated Gαs conformational ensembles at equilibrium, one consisted of particularly long-lived, unique conformational states that required minutes to convert to the crystallographically known GTPγS-bound Gαs conformation. The observed slow closing of the Gαs AHD after GTP binding was unexpected, and confirmation of the existence of long-lived intermediate states in a cellular context will be potentially required using single-molecule imaging techniques. Nevertheless, our finding is of interest when considering the possible effects of long-lived intermediate states on the molecular function of Gs, such as activation cycle kinetics, downstream signaling, or GTPase activity.

As the Gαs AHD closing is a very slow process, we hypothesized the existence of a molecule that regulates the closing kinetics by binding to the Gαs AHD, and identified MAGE D2 as a Gαs AHD-binding partner that accelerates the AHD closing kinetics (Figure 5, step iii-b). The precise mechanism by which MAGE D2 interacts with the long-lived intermediate states of Gαs to shorten their lifetime or accelerate AHD closing is unknown but worth investigating, possibly via high-resolution structural determination of the Gαs–MAGE D2 complex.

Gα binds to various proteins, including GEFs (e.g., GPCRs, Ric-8A, GIV/Girdin, or PLCδ4b), GAPs (e.g., RGSs), effector proteins (e.g., adenylate cyclases or phospholipase C), and other molecules (e.g., GRKs) (Baltoumas et al., 2013; Ghosh and Garcia-Marcos, 2020; Maziarz et al., 2018; Syrovatkina et al., 2016; Tesmer et al., 1997b). Several high-resolution structures of Gα with its binding protein have been published, which all indicate the RD as the main binding interface for these binding partners. For example, RGSs primarily interact with the Switch regions of the RD (Slep et al., 2001; Soundararajan et al., 2008; Tesmer et al., 1997a), although minor interactions are also seen with the AHD when Gα is the Gαq subtype (Nance et al., 2013), and GEFs and effector proteins almost exclusively interact through the RD (Kalogriopoulos *et al*., 2019; Lutz et al., 2007; McClelland et al., 2020; Qi et al., 2019a; Slep *et al*., 2001; Waldo et al., 2010).

To our knowledge, MAGE D2 is the first protein found to interact primarily with a Gα AHD, specifically the Gαs AHD, and to accelerate its closing, which potentially regulates the G protein activation cycle. Notably, the AHD of different Gα subtypes have relatively low sequence identity (approximately 27–42% among the Gαs, Gai1, Gαq, and Gα12 AHDs) compared with that of their RD cores (without αN; approximately 45–58% identity among the Gαs, Gαi1, Gαq, and Gα12 RD cores) (Figure S7). Thus, it is tempting to speculate that the AHD might contribute to the functional selectivity of different Gα subtypes via their binding to different AHD-binding proteins. As MAGE D2 is the first identified Gα AHD-binding protein, we here coin the name **G**α **A**HD **B**inders (GABs). Further studies are needed to identify GABs for other Gα subtypes as well as to understand their molecular functions in G protein signaling.

### Limitation of the study

The main limitation of our study is that it does not provide detailed structural information about Gαs dissociation from the receptor or Gβγ. Not only do HDX-MS technical limitations prevent us from distinguishing the sequence of conformational changes occurring before 5 s of GTPγS addition, but enhanced sampling by metadynamics simulations is only focused on the exploration of the separation between AHD and RD. An additional caveat is that the timeline monitored in the *in vitro* experimental studies (5 s to 30 min) may be different from the timeline required for Gα activation in a cellular context.

## SUPPLEMENTAL INFORMATION

Supplemental information includes eight figures, three tables, and one supplementary data and can be found with this article online.

## ACKNOWLEDGEMENTS

Computations were supported through the computational resources and staff expertise provided by Scientific Computing at the Icahn School of Medicine at Mount Sinai, and were run on resources available through the Office of Research Infrastructure of the National Institutes of Health under award number S10OD026880. The content is solely the responsibility of the authors and does not necessarily represent the official views of the National Institutes of Health. This work was supported by grants from the National Institutes of Health (DA045473 to M.F.) and from the National Research Foundation of Korea funded by the Korean government (NRF-2021R1A2C3003518 and NRF-2019R1A5A2027340 to K.Y.C., and NRF-2019R1C1C1010675 to J.L.).

## AUTHOR CONTRIBUTIONS

D.A. analyzed HDX-MS data, developed and performed the TrIQ used for measuring the AHD movement, prepared MAGE D2 MHD and Gαs AHD, and performed MST analysis. D.P. developed the novel Bayesian integrative modeling framework and used it to derive the proposed kinetic model of Gαs activation upon GTPγS binding. N.M.D. prepared GPCR–Gs complexes, performed HDX-MS experiments, and analyzed the mass spectrometry data. J.X. prepared β_2_AR for TrIQ. L.S.E. carried out the metadynamics simulations of the nucleotide-free β_2_AR–Gs complex, and clustered the simulation results. A.S. carried out the metadynamics simulations of the GTPγS-bound crystal structure. M.W.Y. prepared constructs for Y2H library screening. J.K., D.G., and J.L. carried out the immunofluorescence and immunoprecipitation assays. Y.D. prepared β_2_AR and Gs for HDX-MS. M.F. and K.Y.C. initiated the project, supervised the research, analyzed the data, and wrote the manuscript with contributions from all authors.

## INCLUSION AND DIVERSITY

We support inclusive, diverse, and equitable conduct of research.

## STAR METHODS

### RESOURCE AVAILABILITY

#### Lead contact

Further information and requests for resources and reagents should be directed to and will be fulfilled by the lead contact, Ka Young Chung

#### Materials availability

All unique/stable reagents generated in this study are available from the lead contact with a completed materials transfer agreement.

#### Data and code availability

##### Data

Data reported in this paper will be shared by the lead contact upon request.

##### Code

All original code has been deposited at zenodo and is publicly available as of the date of publication. The DOI is listed in the key resources table.

Any additional information required to reanalyze the data reported in this paper is available from the lead contact upon request.

### EXPERIMENTAL MODEL AND SUBJECT DETAILS

Human β_2_AR was expressed in Sf9 insect cells infected with BestBac recombinant baculovirus (Expression Systems, Davis, CA, USA). The Gs heterotrimer for HDX-MS was expressed in *Trichoplusia ni* insect cells (Expression Systems). Wild-type or mutant Gαs for TrIQ was expressed in the *Escherichia coli* BL21 (DE3) strain, and Gαs AHD and MAGE D2 MHD were expressed in *E. coli* LOBSTR strain.

### METHODS DETAILS

#### Expression and purification of the β_2_AR

The full-length β_2_AR was prepared as previously described (Du *et al*., 2019). In brief, recombinant baculovirus was prepared using the Bestbac expression system (Expression Systems) and pVL1392 as a transfer vector. The full-length β_2_AR was expressed by infecting Sf9 cells with second-passage baculovirus stock. The β_2_AR antagonist alprenolol (2 μM) was added to stabilize the receptor during expression. The infected cells were harvested after 48 h of incubation at 27°C. Cell pellets were lysed, and the receptor was extracted from the cell membrane using a Dounce homogenizer in solubilization buffer [20 mM HEPES, pH 7.5, 100 mM NaCl, 1% dodecylmaltoside (DDM), 1 μM alprenolol, 2.5 μg/mL leupeptin, and 160 μg/mL benzamidine] for 1 h at room temperature (22–25°C). The receptor bearing the N-terminal FLAG tag was then captured by M1 antibody affinity chromatography (Sigma-Aldrich, St. Louis, MO, USA) and further purified by affinity chromatography using alprenolol–Sepharose as previously described (Rosenbaum et al., 2007) to select functional receptors. The purified receptor with alprenolol was tandem-linked to the M1 FLAG column and washed with HMS–CHS buffer (20 mM HEPES, pH 7.5, 350 mM NaCl, 0.1% DDM, 0.01% cholesterol hemisuccinate) for removal of alprenolol to obtain an unliganded receptor. The receptor was then eluted from the M1 resin with HMS–CHS buffer supplemented with 5 mM EDTA, 200 μg/mL free FLAG peptide, and 10 μM BI-167107. Size-exclusion chromatography with a Superdex-200 column (GE Healthcare Life Science, Uppsala, Sweden) equilibrated in HLS–CHS buffer (20 mM HEPES pH 7.5, 150 mM NaCl, 0.1% DDM, 0.01% cholesterol hemisuccinate, 2 μM BI-167107) was finally used to polish the receptor. BI-167107 is a high-affinity β_2_AR agonist that was used to solve the high-resolution crystal structure of β_2_AR (Rasmussen *et al*., 2011). The receptor was concentrated to 150 μM for preparing samples for HDX-MS analyses. The purity of the sample was higher than 95%, as assessed by sodium dodecyl sulfatepolyacrylamide gel electrophoresis (SDS-PAGE).

#### Expression and purification of Gs and Gβγ

Heterotrimer Gs and Gβγ were prepared as previously described (Du *et al*., 2019). In brief, for heterotrimeric Gs purification, bovine Gαs (short isoform), His6-rat Gβ_1_, and bovine Gγ_2_ were co-expressed in *Trichoplusia ni* insect cells grown in ESF 921 medium (Expression Systems) using two separate *Autographa californica* nuclear polyhedrosis viruses containing the Gαs and Gβγ genes. Cell pellets expressing G protein subunits were resuspended in lysis buffer (10 mM Tris, pH 7.4, 100 μM MgCl_2_, 5 mM β-ME, 10 μM GDP, 2.5 μg/mL leupeptin, and 160 μg/mL benzamidine) for 30 min at room temperature (22–25°C). The lysate was spun for 15 min at 18,000 *g*, and then the pellet was homogenized in solubilization buffer (20 mM HEPES, pH 7.5, 100 mM NaCl, 1% sodium cholate, 0.05% DDM, 5 mM MgCl_2_, 2 μL calf intestinal alkaline phosphatase, 5 mM β-ME, 10 μM GDP, 5 mM imidazole, 2.5 μg/mL leupeptin, and 160 μg/mL benzamidine) using a 100-mL Dounce homogenizer and tight pestle. The heterotrimeric Gs was purified using nickel-bound Chelating Sepharose Fast Flow and Q Sepharose resin. The purified protein was passed through a 0.22-μm filter and spin-concentrated in a 10-kDa MWCO concentrator (Millipore, Burlington, MA, USA) to approximately 20 mg/mL. The methods used for the expression and purification of lipidated Gβ_1_γ_2_ were very similar to those described for the Gs heterotrimer, except that GDP and MgCl_2_ were removed from the buffer formulation.

#### Expression and purification of Gαs, Gαs AHD, and MAGE D2 MHD

Recombinant Gαs containing an N-terminal His-tag and HRV 3C protease cleavage site was constructed in the pET28a vector. The cDNA of the human MAGE D2 MHD sequence (GenBank: NM_014599; amino acids 261–521) was synthesized and inserted into pET-28a with an N-terminal 6 histidine tag-TEV cleavage site. The cDNA of the human Gαs AHD (GenBank: NM_080426; amino acids 68–206) was synthesized and inserted into pET-28a with an N-terminal 6 histidine tag-thrombin cleavage site and C-terminal FLAG tag. Gαs was transformed into *E. coli* BL21 (DE3), and MAGE D2 MHD and Gαs AHD were transformed into *E. coli* LOBSTR for protein expression. The cells were grown in Terrific Broth in the presence of antibiotic at 37°C until the optical density at 600 nm reached 0.6–0.8. Gαs or MAGE D2 MHD expression was induced by adding 0.5 mM isopropyl β-D-1-thiogalactopyranoside (IPTG) following incubation at 25°C for 18 h or 4 h, respectively. Gαs AHD expression was induced by adding 0.5 mM IPTG following incubation at 37°C for 4 h. For protein purification, the bacterial pellets were suspended in lysis buffer [20 mM HEPES, pH 7.4, 300 mM NaCl, 5 mg/mL lysozyme, 1:1000 protease inhibitor cocktail, 2.5 μg/mL leupeptin, 10 μg/mL benzamidine, 100 μM tris(2-carboxyethyl)phosphine (TCEP)] and incubated for 30 min. The lysates were spun down at 18,000 *g* for 20 min at 4°C, and the supernatant was collected and loaded onto an Ni-NTA column pre-equilibrated with the lysis buffer supplemented with 20 mM imidazole. The resin was washed with wash buffer (20 mM HEPES, pH 7.4, 100 mM NaCl, 100 μM TCEP, 20 mM imidazole), and the bound proteins were eluted with elution buffer (20 mM HEPES, pH 7.4, 100 mM NaCl, 100 μM TCEP, 200 mM imidazole). The eluted products were loaded onto a Superdex-200 (10/300) column equipped with an ÄKTA FPLC system (GE Healthcare Life Sciences), and the purified proteins were eluted with a second elution buffer (20 mM HEPES, pH 7.4, 100 mM NaCl, 100 μM TCEP). For Gαs, before loading onto the Superdex-200 (10/300) column, the His-tag was removed by incubation with 3C protease at 4°C overnight. Through the purification step, for Gαs purification, 2 mM MgCl_2_ and 20 μM GDP were supplemented in the lysis, wash, and elution buffers.

#### Nucleotide-free β_2_AR–Gs complex formation

The nucleotide-free β_2_AR–Gs complex was prepared as previously described (Du *et al*.,2019). In brief, 65 μM of Gs was mixed with 65 μM β_2_AR at room temperature (22–25°C). Apyrase (200 mU/mL) was added after 90 min of incubation to hydrolyze the GDP released from Gαs, and incubation was proceeded for a further 90 min.

#### Pulse-labeling HDX-MS

For pulse-labeling deuterium exchange, the β_2_AR, GDP-bound Gs, and nucleotide-free β2AR–Gs complex were prepared at a concentration of 65 μM. Four microliters of the β_2_AR or GDP-bound Gs was mixed with 26 μL of D_2_O buffer (20 mM HEPES, pD 7.5, 100 mM NaCl, 2 μM agonist, 100 μM TCEP, 0.1% DDM in D_2_O) and incubated for 10 s at room temperature (22–25°C). Four microliters of the nucleotide-free β2AR–Gs complex was aliquoted at specific time points (before adding GTPγS and after adding 500 μM GTPγS for 5 s, 30 s, 2 min, 5 min, 10 min, and 30 min) at room temperature (22–25°C) and mixed with 26 μL of D_2_O buffer at room temperature (22–25°C). All deuterium-exchanged samples were quenched by 30 μL of ice-cold quench buffer (0.1 M NaH_2_PO_4_, 20 mM TCEP, pH 2.01), immediately frozen on dry ice, and stored at – 80°C. For non-deuterated samples, 4 μL of protein samples (65 μM) were mixed with 26 μL of H_2_O buffer (20 mM HEPES, pH 7.4, 100 mM NaCl, 0.1% DDM in H_2_O) to which 30 μL of ice-cold quench buffer was added and snap-frozen on dry ice.

The quenched samples were digested and analyzed by an HDX-ultra-pressure liquid chromatography (UPLC)-electrospray ionization (ESI)-MS system (Waters, Milford, MA, USA) as previously described (Du *et al*., 2019). In brief, quenched samples were thawed and immediately injected into an immobilized pepsin column (2.1 × 30 mm) (Life Technologies, Carlsbad, CA, USA) at 10°C. Peptide fragments were subsequently collected on a C18 VanGuard trap column (1.7 μm × 30 mm) (Waters, Milford, MA, USA) for desalting and then isolated by UPLC using an Acquity UPLC C18 column (1.7 μm, 1.0 × 100 mm) (Waters). To minimize the back-exchange of deuterium to hydrogen, the system, including the trapping column and UPLC column, was maintained at 0.5°C during the analysis, and all buffers were adjusted to pH 2.5. Mass spectral analyses were performed with a Xevo G2 quadruple time-of-flight system equipped with a standard ESI source in MS^E^ mode (Waters) and positive-ion mode. All settings/conditions for the system were as previously reported (Du *et al*., 2019). Peptic peptides were identified in non-deuterated samples with ProteinLynx Global Server 2.4 (Waters) with variable methionine oxidation modification, and the peptides were filtered according to a peptide score of 6. To process the HDX-MS data, the amount of deuterium in each peptide was determined by measuring the centroid of the isotopic distribution using the DynamX 2.0 software package (Waters). The average back-exchange level in our system was 30–50%, which was not corrected because the β_2_AR only aggregated in the fully deuterated sample condition, and the analyses were based on a comparison of different states of proteins.

#### Measurement of the separation between the RD and AHD

To measure the separation between the RD and AHD, we used a tryptophan-induced bimane quenching (TrlQ) method. First, we mutated residue A303 in the Gαs RD, which is adjacent to the AHD, into Cys for monobromobimane labeling. Residue E164 in the Gαs AHD was mutated into Trp. The distance between A303 and E164 is 10.7–10.8 Å in the GDP-bound or GTPγS-bound Gαs crystal structures (PDB: 6EG8 and PDB: 1AZT, respectively). At this distance, a Trp residue at position 164 would be close enough to quench the fluorescence of the bimane at position 303. In the β_2_AR–Gs crystal structure (PDB: 3SN6), these residues are further apart due to displacement of the AHD, and therefore the quenching effect disappears.

Solvent-exposed Cys residues (C174, C200, C237) in Gαs were mutated into Ser to prevent unwanted bimane labeling. All mutations were generated by site-directed mutagenesis using polymerase chain reaction (PCR). The engineered Gαs was incubated with a 2-molar excess of monobromobimane at room temperature (22–25°C) for 1 h. The unbound bimane was removed by buffer exchange with the reaction buffer (20 mM HEPES, pH 7.4, 100 mM NaCl, 2 mM MgCl_2_, 10 μM GDP, 0.1% DDM, 0.01% CHS, 1 μM BI-167107). The bimane-labeled Gαs formed a heterotrimer with Gβγ by incubating at room temperature (22–25°C) for 30 min. To form the β_2_AR–Gs complex, the heterotrimeric G protein was mixed with the β_2_AR for 90 min, followed by another 90 min incubation with Apyrase, which hydrolyzes GDP. The bimane fluorescence was measured by excitation at 390 nm (bandwidth 10 nm) and emission at 485 nm (bandwidth 10 nm) using a BioTek Synergy 2 microplate reader (Santa Clara, CA, USA). GTPγS (500 μM) was added, and the bimane fluorescence was measured every 10 s for a total of 2,400 s. All experiments were conducted at room temperature (22–25°C).

#### Nucleotide exchange assay

The nucleotide exchange function of Gαs was evaluated by the binding of BODIPY-FL-GTPγS (Invitrogen, Waltham, MA, USA) into the GDP-bound Gαs. BODIPY-FL-GTPγS (250 nM) was prepared in an imaging buffer (20 mM Tris pH 8.0, 1 mM EDTA, 10 mM MgCl_2_, 100 μM TCEP). The fluorescence baseline was recorded by TriStar2 S LB 942 Multimode Microplate Reader (Berthold Technologies, Bad Wildbad, Germany) for 120 s. The samples were excited at 485 nm (bandwidth, 14 nm) and detected the emission at 535 nm (bandwidth, 25 nm) in a 96-well black plate. 1.5 μM of GDP-bound Gαs was added, and the fluorescence was measured every 10 s for a total of 2,400 s. All experiments were conducted at room temperature (22–25°C).

#### Y2H assay

Y2H screening was conducted by Panbionet (Pohang, South Korea). The AHD (residues 68–192) of *GNAS* cDNA (375 bp) was amplified by PCR. The PCR product was cloned into a pGBKT7 vector containing the DNA-binding domain (BD) of GAL4. *Saccharomyces cerevisiae* (yeast) strain PBN240 (Panbionet) was co-transformed with the GAL4 DNA-BD fused to GNAS68–192 and the cDNA activation domain (AD) library. The PBN204 strain contains three reporters (*URA3, lacZ*, and *ADE2*) that are under the control of different *GAL* promoters. The protein–protein interaction was tested using three independent reporters with different types of GAL4-binding sites to reduce the possibility of picking up false-positive candidates; in some cases, reporter gene expression can be activated by an AD fusion protein in a particular *GAL* promoter-specific manner. Yeast transformants of the *GNAS* bait and the human kidney cDNA AD library were spread on a selection medium [SD-leucine, tryptophan, uracil (SD-LWU)] that supports the growth of yeasts with bait and prey plasmids, yielding proteins interacting with each other. After selecting yeast colonies on uracil-deficient media, the activity of beta-galactosidase was monitored. The URA+ and lacZ+ colonies were monitored to see if they were able to grow on adenosine-deficient media. To confirm the interaction, the prey parts of DNA from 60 U+A+Z+ candidates were amplified by PCR or by *E. coli* transformation, and then the amplified candidate prey was reintroduced into yeast with the *GNAS* bait plasmid or with a negative control plasmid.

#### Microscale thermophoresis (MST) assay

The MAGE D2 MHD was fluorescently labeled using Monolith Protein Labeling Kit RED-NHS 2nd Generation (NanoTemper, München, Germany) according to the manufacturer’s instructions. Labeled MAGE D2 MHD (50 nM) and two-fold serially diluted Gαs AHD (7.629 nM–25 μM) were mixed in the interaction buffer (20 mM HEPES, pH 7.4, 100 mM NaCl, 100 μM TCEP, 0.05% Tween-20). The measurement was conducted with Monolith NT.115 (NanoTemper) at 20% excitation power and medium MST power at 25°C. The results were acquired from MST-on of 5 s.

#### Immunostaining

cDNAs encoding the AHD open reading frames of Gαs cDNA were synthesized according to the GenBank ID NM_080426 and were cloned into the AsiSI and PmeI restriction sites of pCMV6-AC-GFP (Origene, Rockville, MD, USA) using the Gibson assembly (NEB, Ipswich, MA, USA). The 3X FLAG-tag sequence (5-gactacaaagaccatgacggtgattataaagatcatgacatcgattacaaggatgacgatgacaag-3’) was inserted before the stop codon of the Gαs AHD. HEK293T cells were seeded 24 h prior to transfection at a density of 2 × 10□ cells onto cover glasses in a 24-well plate. pCMV6-Gαs AHD-3XFLAG was prepared in Transfection Optimization Media (TOM) and transfected into cells using Lipofectamine 3000. The total amount of plasmids was adjusted to 400 ng. After 24 h, the cells were washed with phosphate-buffered saline (PBS) and fixed for 15 min using 4% of formaldehyde in PBS. Fixed cells were permeabilized for 15 min with 0.1% Triton X-100. Cells were then blocked with 1.5% of bovine serum albumin in PBS for 60 min and labeled with the indicated antibodies overnight. Alexa Fluor-488 and −647 antibodies were used as secondary antibodies. Sixty minutes after treatment with secondary antibodies, the cells were treated with Hoechst 33342 solution (Invitrogen) for 20 min. After the last wash, the cells were dried and mounted with mounting solution.

#### Immunoprecipitation and immunoblot analysis

HEK293T cells were seeded 24 h prior to transfection at a density of 5 × 10□ cells in 100-mm dishes. pCMV6-MAGE D2-turboGFP and pCMV6-Gαs AHD-3XFLAG were prepared in TOM and transfected into cells using Lipofectamine 3000. The total amount of plasmids was adjusted at 1000 ng. After 24 h, the cells were washed with PBS and lysed with RIPA buffer in the presence of a protease inhibitor cocktail. SureBeads Protein G magnetic beads (Bio-Rad, Hercules, CA, USA) and anti-FLAG (1:1000) were added to the transferred lysates and incubated overnight. Proteins bound to antibodies and protein G Sepharose were collected by centrifugation and washed three times with RIPA buffer. The precipitated samples were boiled in SDS-PAGE sample loading buffer (Abpbio, Beltsville, MD, USA) for 10 min at 95°C. Proteins and a molecular-weight marker (Bio-Rad) were subjected to Tris-glycine SDS-PAGE and then transferred onto polyvinylidene fluoride membranes. The membrane was blocked with 5% skim milk in TBS-T buffer, incubated with the primary antibody at 4°C overnight, and then incubated with horseradish peroxidase-conjugated secondary antibody for 60 min at room temperature (22–25°C). Immunoblots were visualized using AbSignal western blotting detection reagent (AbClon, Seoul, Republic of Korea) and detected using Bio-Rad ChemiDoc (Bio-Rad).

#### Systems setup for simulations

The crystal structure of the activated bovine Gαs bound to GTPγS and Mg^2+^ (PDB ID: 1AZT (Sunahara *et al*., 1997)) was used as a reference conformation of the protein with a closed AHD. All non-protein atoms except for the Mg^2+^ ion and GTPγS were removed. Missing residues 9–34 and 392–394 were modeled based on the crystal structure of human Gs in complex with GDP (PDB ID: 6EG8; (Liu *et al*., 2019)), using the “Protein Splicing” tool within the Schrödinger’s Maestro software package (Schrodinger Release 2019-4, 2019). Residues 70–73, which were missing from both crystal structures, were added using the “Crosslink Proteins” panel in Maestro, and their conformation was predicted using the “Loop lookup” option to search for known geometries in the PDB. The nucleotide-free form of Gαs bound to Gβγ subunits and the agonist BI167107-bound β_2_AR (PDB: 3SN6 (Rasmussen *et al*., 2011)) were also prepared using standard protocols of the Maestro software package (Schrodinger Release 2019-4, 2019), with engineered residues in the 3SN6 construct converted into wild-type human residues. Missing short segments in β_2_AR (176–178) or Gαs (53–64, 70–87, and 256–262) were built either with MODELLER 9.19 (Marti-Renom et al., 2000; Webb and Sali, 2016) based on available templates (PDBs: 3P0G (Rasmussen *et al*., 2011), 1AZS (Tesmer *et al*., 1997b), 6EG8 (Liu *et al*., 2019), and 1AZT (Sunahara *et al*., 1997), respectively) or *ab initio* (Gαs 65–69 and 203–204), whereas longer missing segments or termini (β_2_AR 1–29, 240–264, and 342–413, as well as Gαs 1–8 and *Gγ* 1–4 and 63–71) were not modeled and their ends were capped.

#### Molecular dynamics simulations for energy minimization and equilibration

After adding hydrogens and Mg^2+^ in the nucleotide binding pocket, the aforementioned GTPγS-bound Gαs system with GTPγS modeled using the CHARMM general force field (CGenFF) (Vanommeslaeghe et al., 2010) was immersed in a dodecahedron water box of TIP3P water molecules (Jorgensen et al., 1983) in such a way that there were at least 10 Å of water molecules between the protein and the boundary of the unit cell in all directions. The nucleotide-free β_2_AR-Gs system with CGenFF-derived parameters for the β_2_AR ligand BI-167107 was embedded in a POPC/10% cholesterol membrane patch and solvated in TIP3P water to form a 122 × 122 × 170 Å^3^ simulation box. The NaCl concentration was set to 0.15 M, and the proteins were modeled using the CHARMM36m (Huang et al., 2017) force field, with a 12 Å cutoff set for non-bonded interactions. Long-range electrostatic interactions were calculated using the particle-mesh Ewald method (Darden et al., 1993). All bonds containing hydrogen atoms were fixed using the LINCS algorithm (Hess et al., 1997), while hydrogen masses were increased to 4 amu using the mass repartition scheme (Feenstra et al., 1999), so that production molecular dynamics simulations could be executed with a time step of 4 fs. Both GTPγS-bound Gαs and nucleotide-free β_2_AR-Gs systems were minimized and subsequently equilibrated to a temperature of 300 K using the Nose-Hoover thermostat (Hoover, 1985; Nose, 1984) and to a pressure of 1 atm using the Parinello-Rahman barostat (Nose and Klein, 1983; Parrinello and Rahman, 1981) by 50 ns and 10 ns, respectively, unrestrained molecular dynamics simulations using the Gromacs 2021.4 software package (Abraham et al., 2015).

#### Metadynamics simulations for exhaustive conformational sampling

The conformational rearrangement between the closed and open states of the Gαs AHD in both GTPγS-bound Gαs and nucleotide-free β_2_AR–Gs systems was simulated using well-tempered metadynamics simulations (Barducci *et al*., 2008; Laio and Gervasio, 2008; Laio and Parrinello, 2002) to enhance sampling along two collective variables. Specifically, CV1 corresponded to a plane projection of the angle between the two vectors connecting the Cα atoms of the AHD residues 95 and 106 or the RD residues 58 and 341 of the Gαs subunits, whereas CV2 accounted for the deviation of the contact map between the two domains with respect to one derived from PDB: 1AZT. This deviation was measured by monitoring the distances between the Cα atoms of all pairs of residues on the two domains that are within 8 Å of each other in the starting (closed) structure of 1AZT. Smooth switching between contacts being formed and broken was accomplished using a rational switching function (1 – (*r*/*r*_0_)^6^)/(1 – (*r*/*r*_0_)^12^), where *r* is the distance between the Cα atoms of the interacting residues and r_0_ was set to 8.0 Å. Well-tempered metadynamics simulations were performed with Gromacs 2021.4 interfaced with the PLUMED library (Bonomi et al., 2009; Tribello et al., 2014), adding Gaussian hills every 1000 steps with an initial height of 1.0 kJ/mol and a bias factor of 10, and with a width of 0.05 rad and 0.1 for the angle and contact dimensions, respectively. Reweighting factors to recover unbiased values from the simulations were calculated using an established methodology (Tiwary and Parrinello, 2015). Simulations were carried out at a temperature of 300 K and pressure of 1 atm with a time step of 4 fs, as described above for the unbiased simulations. Data were recorded every 20 ps. The total metadynamics simulation time was 7.6 μs and 7.0 μs for the GTPγS-bound Gαs and nucleotide-free β_2_AR–Gs systems, respectively. Convergence was assessed by plotting changes in free energy as a function of the two aforementioned CVs, using first-quarter, half, three-quarters, and the whole trajectories (Figure S4B).

#### Clustering of metadynamics-derived conformational states

To identify the different conformations sampled by the GTPγS-bound Gαs and nucleotide-free β_2_AR–Gs systems during metadynamics simulations, the resulting trajectories were clustered using the pairwise distances between every third Cα atom in Gαs (6,441 distance pairs). Residues 9–32 of the αN helical domain and residues 33– 38 of the αN/β1 loop were left out from this analysis because they were very flexible during simulations of GTPγS-bound Gαs, which was assumed to be due to the absence of Gβγ. The pairwise distances were calculated using batches of 20 frames from each trajectory (18,910 and 17,601 frames for the GTPγS-bound Gαs and nucleotide-free β_2_AR–Gs systems, respectively), and their values were projected onto the first 10 time-structure independent component analysis (tICA) components. Pooled tICA features from the two systems were partitioned in 10 groups using k-means clustering, and these groups were further split into 16 structural clusters, *R_s_*, with nine pertaining to the GTPγS-bound Gαs system and seven to the nucleotide-free β_2_AR–Gs system. Finally, 10 frames from each cluster were sampled as representatives, which were used to calculate the values of the protection factors 〈*PF_j_*〉_*s*_ and deuterium uptake *D_J_*(*s*) as input for the kinetic modeling described below. Calculation of the RMSD values, as well as the atom–atom distances needed for the estimation of the protection factors (see below), was carried out with MDTraj (McGibbon et al., 2015), whereas the tICA and clustering analyses were performed with PyEMMA (Scherer et al., 2015).

#### Prediction of protection factors from metadynamics-sampled conformations

HDX experiments probe the rate of exchange of backbone amide hydrogens with deuterium (^2^H or D) upon solvation into D_2_O. The hydrogen/deuterium exchange rate of a specific residue is affected by the macroscopic environment (e.g., pressure, temperature, ionic strength of the solvent), as well as the nature of the amino acids immediately preceding and following the residue in the protein sequence (Bai et al., 1993; Molday et al., 1972), long-range electrostatic interactions (Kim and Baldwin, 1982), and local distribution of charged residues (Wu et al., 2009). In particular, the involvement of backbone amide hydrogens in hydrogen bonding with buried residues of a protein can slow down this exchange. This slowdown is commonly expressed in terms of protection factors *P* = *k*_obs_/*k*_int_, where *k*_obs_ is the observed exchange rate and *k*_int_ is the “intrinsic” exchange rate in the fully unprotected, solvent-accessible state. Values of *k*_int_ for each residue in a protein are typically not available and are instead approximated by the values measured for short peptides flanked by poly-D/L-alanine. In this work, we adopted intrinsic rates (Bai *et al*., 1993) of the following form:

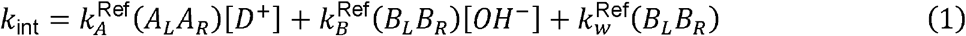

where [*D*^+^] and [*OH*^-^] are the deuteron and hydroxide concentrations, respectively; 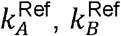, and 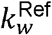 are values for side chain-specific acid, base, and water rate constants, respectively; and *A_L_, A_R_, B_L_*, and *B_R_* are correction factors that depend on the residue on the left and right side of the reference residue, respectively.

A semi-empirical approximation of protection factors of a protein in a given conformation *R* introduced by Vendruscolo et al. (Vendruscolo et al., 2003), and later generalized by Wan et al. (Wan et al., 2020), expresses the protection factors for the protein in a conformational ensemble *R_s_* as:

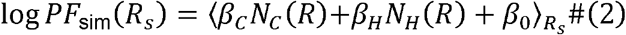

Where *N_C_*(*R*) is the number of heavy atoms within a given cut-off distance *d_C_* from the amide nitrogen, *N_H_*(*R*) is the number of hydrogen bonds formed by the amide, the *βs* are model coefficients, and the average is calculated over all conformations *R* in the ensemble *R_s_*. To calculate, using eq. (2), the protection factors in the clusters *R_s_* identified for our systems, we defined *N_c_* and *N_H_* using sigmoid switching functions:

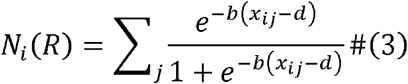

where *x_ij_* is the distance between the amide on residue *i* and atom *j* in conformation *R*, and *b* and *d* are parameters that determine the effective cut-off distance used to derive the values of *N_c_* and *N_H_*. The sum is extended to all heavy atoms *j* for the calculation of *N_c_*, and to all hydrogen bond acceptors for the calculation of *N_H_*. The original application (Vendruscolo *et al*., 2003) used *β_0_* = 0, and the coefficients *β_c_* and *β_H_* were obtained by the least-squares fit of eq. 2 to the HDX experimental results obtained for the native state of seven different globular proteins, and fixed to *β_c_* ≈ 0.35 and *β_H_* ≈ 2. The values of the distance cutoffs were fixed to *d_c_* = 6.5 Å, and a cutoff of *d_H_* = 2.4 Å between hydrogen donors and acceptors was used to identify hydrogen bonds (Vendruscolo *et al*., 2003).

Bayesian inference was used to determine the parameters *{β_C_, β_H_, β_0_, b, d_C_, d_H_*} = *θ* of the model as proposed by Wan et al. (Wan *et al*., 2020). Specifically, the joint distribution of the parameters was expressed as:

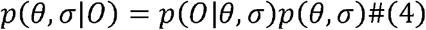

where *p*(*O*|*θ*, *σ*) is the likelihood of the observed data *O* derived from eq. (2) with a Gaussian error structure of variance *σ*^2^:

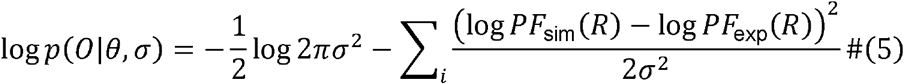

The full posterior *p*(*θ*, *σ*|*O*) in the original implementation was sampled using experimental data available on two globular proteins (Vendruscolo *et al*., 2003). In this work, we adopted the distribution sampled in Wan et al. (Wan *et al*., 2020) as the prior distribution for HDX modeling.

#### Prediction of quenching data using metadynamics-sampled conformations

The separation and relative orientation between the Gαs RD and ADH were probed experimentally by measuring the quenching of the emission intensity of bimane, an organic fluorophore, by electron-donating residues (e.g., tryptophan and tyrosine) or moieties (e.g., guanine), using photoinduced electron transfer (PET) (Callis, 2014; Mansoor et al., 2010; Mansoor et al., 2002). Unlike Förster resonance energy transfer, which typically monitors longer distances, PET measures quenched fluorescence upon van der Waals contact, thus providing a better estimate of molecular contacts (Jones Brunette and Farrens, 2014). Given a conformational ensemble characterized by the probability distribution μ(*R*), we express the bimane fluorescence intensity as:

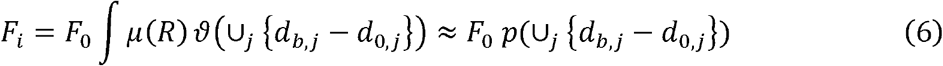

where *F_i_* is the measured quenched fluorescence, *F*_0_ is the bimane intensity in the absence of quenching, *d_b,j_* is the distance between the bimane and electron-donating moiety *j*, *d*_0,*j*_ is the appropriate distance cutoff for the quenching moiety, and ∪_*j*_[*d_b,j_* – *d_0,j_*] represents the case wherein at least one quencher moiety is close enough to the fluorophore. Adopting previously published reference values (Mansoor *et al*., 2002), we used 15 Å as an appropriate distance cutoff for quenching induced by tryptophan and 10 Å for quenching induced by tyrosine.

#### Bayesian inference to update metadynamics-derived prior probabilities of conformational states with pulsed HDX-MS and fluorescence quenching data

Pulsed HDX-MS probes the time-dependence of the total deuterium uptake by a protein exposed to D_2_O for short pulses of time. The fraction of deuterated amino acids *D_J_*(*s*) for a peptide segment *J* of the protein in each conformational ensemble *s* can be calculated for a pulse *τ* using the protection factors of the individual residues *j* ∈ *J* of the peptide as:

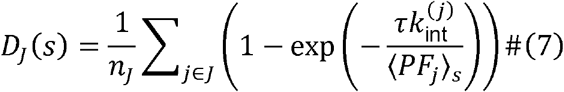

where 〈*PF_j_*〉_s_ is the average protection factor for residue *j* in state *s* and 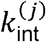 is the intrinsic rate for residue *j*. We assume that the state of the system can be described as a Markov process between conformational states *s*, with a transition rate matrix ***k***, so that the transition probabilities from one metadynamics-derived cluster to another can be expressed as exp(***k**t*). Given this model, the probability that the system is in each state at time t can be expressed as ***π***(*t*) = ***π***(0) exp(***k**t*), where *_π_*(0) is the initial probability distribution and ***k*** is the transition matrix describing the kinetics of the protein. Using these probabilities, the time-dependence of the observed *D_J_* values can be expressed as:

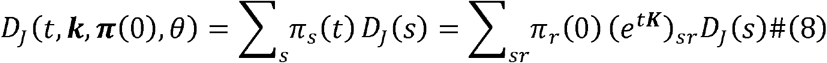

where *θ* is the parameter that determines the values of *D_j_*(*s*) in eq. (7). The loglikelihood of the observed HDX data for a given value of the model parameters (***k,π***(0), *θ*) is

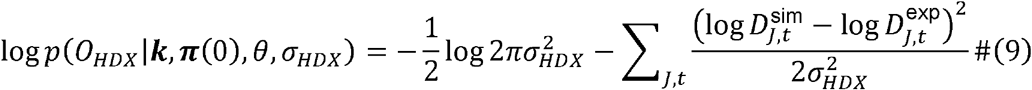

where 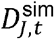 is the deuterated fraction for peptide segment *J* of the protein given by eq. (8), 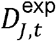 is the corresponding experimental value, and *σ_HDX_* quantifies the uncertainty of experimental measures. The results presented in this analysis were obtained using data for 13 of the peptides reported in Figures 1 and 2 (46–59, p-loop-α1; 108–118, N-terminal region of αA/αB loop; 119–126, αA/αB loop-αB; 126–132, αB; 178–193, αE; 190–197, αE/αF loop-αF; 201–208, Switch I; 256–272, Switch III-α3; 290–297, β5-αG; 297–312, αG-αG/α4 loop; 367–371, β6/α5-N-terminus of α5; 372–381, N-terminal half of α5; and 382–390, C-terminal half of α5).

Similarly, we included the values of the bimane fluorescence quenching in the model inference at the beginning of the experiment and at equilibrium. Following the same approach outlined above for HDX, the ratio of the quenched over non-quenched fluorescence is expressed as the probability that at least one quencher moiety is close enough to the fluorophore

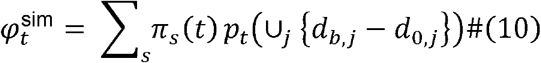

and the likelihood of the observed bimane data is then

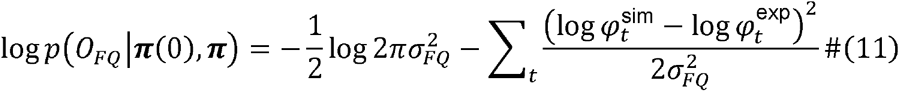

Finally, the posterior joint distribution of the parameters that describe the model can be expressed as:

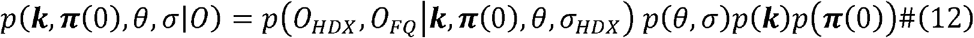

#### Maximum Caliber priors and sampling to derive the transition matrix

To use eq. (12) to determine the transition matrix *k*, the prior distributions *p*(*θ,σ*), *p*(***k***), and *p*(***π***(0)) must first be specified. We used the distributions from Wan et al. (Wan *et al*., 2020) for the model parameters. In line with the experimental setup, we assumed that the prior distribution for the initial probabilities *p*(***π***(0)) is concentrated on states from the nucleotide-free β_2_AR–Gs system. Thus, we used an informative distribution for ***π***(0) by assigning a vanishing probability to the GTPγS-bound Gαs states. We incorporated the maximum entropy (Maximum Caliber) principle (Pressé et al., 2013) into the Bayesian inference by deriving entropic priors *p*(***k***) for the kinetic matrix. Following Caticha and Preuss (Caticha and Preuss, 2004), we express the prior for ***k*** as,

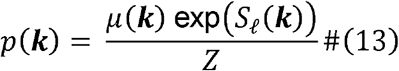

where *Z* = ∫ *μ*(***k***) *e*^S_ℓ_(***k***)^ is a normalization factor and *S_ℓ_*(***k***) is the local relative entropy as a function of the transition matrices ***k*** ∈ ***K***

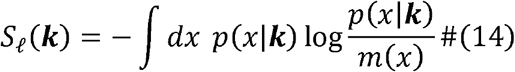

To fix the prefactor μ(***k***) in eq. (13), we chose 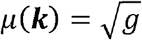, where *g* = det*g_ij_* is the determinant of the information metric induced by *S_ℓ_*(***k***) on the space ***K***:

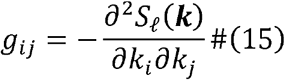

Specifically, if *x* is the space of Markov paths on a discrete set of microstates and ***k*** = {*k_αβ_*} are the rates of transition between the microstates, then the local entropy *S_ℓ_*(***k***) can be written as:

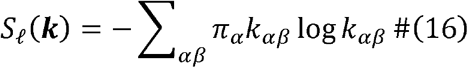

It is convenient to fix the stationary probability *π_α_* and work in the subspace of *K_π_* with that distribution, so that

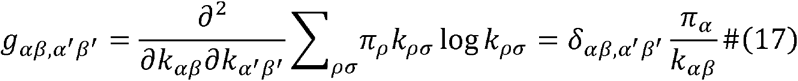

The prior distribution for the transition matrix *p*(***k***|***π***) is therefore

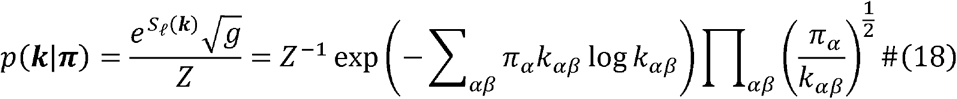

Given that the experimental conditions are such that GTPγS saturates Gαs at equilibrium, we assumed that the prior distribution for the equilibrium probabilities *p*(*π*) is concentrated on states from the GTPγS-bound Gαs simulation and assigned vanishing probabilities to the states sampled by the nucleotide-free β_2_AR–Gs system.

Using this prior and the likelihoods in eq. (9) and (11), we sampled the full posterior distribution *p*(***k***, ***π***(0), *θ*, *σ*|*O*) using a Gibbs sampling algorithm implemented in an in-house Python script. Values of *D_J_*(*t*) and ***π***(*t*) sampled from the posterior distribution (eq. (12)) are reported in Figures S5 and S8, respectively. Predicted populations of the initial state of the systems (at time 0) were obtained from the sampled marginal posteriors of the initial probabilities, the equilibrium distribution of the states (at long time scales) was obtained from Perron eigenvectors of the sampled transition matrices, and the state probabilities at intermediate times (Figure S8) were obtained by direct application of the transition matrix.

### QUANTIFICATION AND STATISTICAL ANALYSIS

For HDX-MS analysis, mass differences bigger than 0.3 Da were considered significant, and a paired *t*-test run in GraphPad Prism software was used to determine the statistically significant differences between individual time points with *p* < 0.05 considered to be statistically significant.

