## Supplementary figures tables for "Gαs slow conformational transition upon GTP binding and a novel Gαs regulator": Suppl Figures and tables_BioRxiv.pdf

G proteins are major signaling partners for G protein-coupled receptors (GPCRs). Although stepwise structural changes during GPCR–G protein complex formation and guanosine diphosphate (GDP) release have been reported, no information is available with regard to guanosine triphosphate (GTP) binding. Here, we used a novel Bayesian integrative modeling framework that combines data from hydrogen-deuterium exchange mass spectrometry, tryptophan-induced fluorescence quenching, and metadynamics simulations to derive a kinetic model and atomic-level characterization of stepwise conformational changes incurred by the $\beta_2$-adrenergic receptor ($\beta_2$AR)-Gs complex after GDP release and GTP binding. Our data suggest rapid GTP binding and GTP-induced dissociation of Gαs from $\beta_2$AR and Gβγ, as opposed to a slow closing of the Gαs α-helical domain (AHD). Yeast-two-hybrid screening using Gαs AHD as bait identified melanoma-associated antigen D2 (MAGE D2) as a novel AHD-binding protein, which was also shown to accelerate the GTP-induced closing of the Gαs AHD.

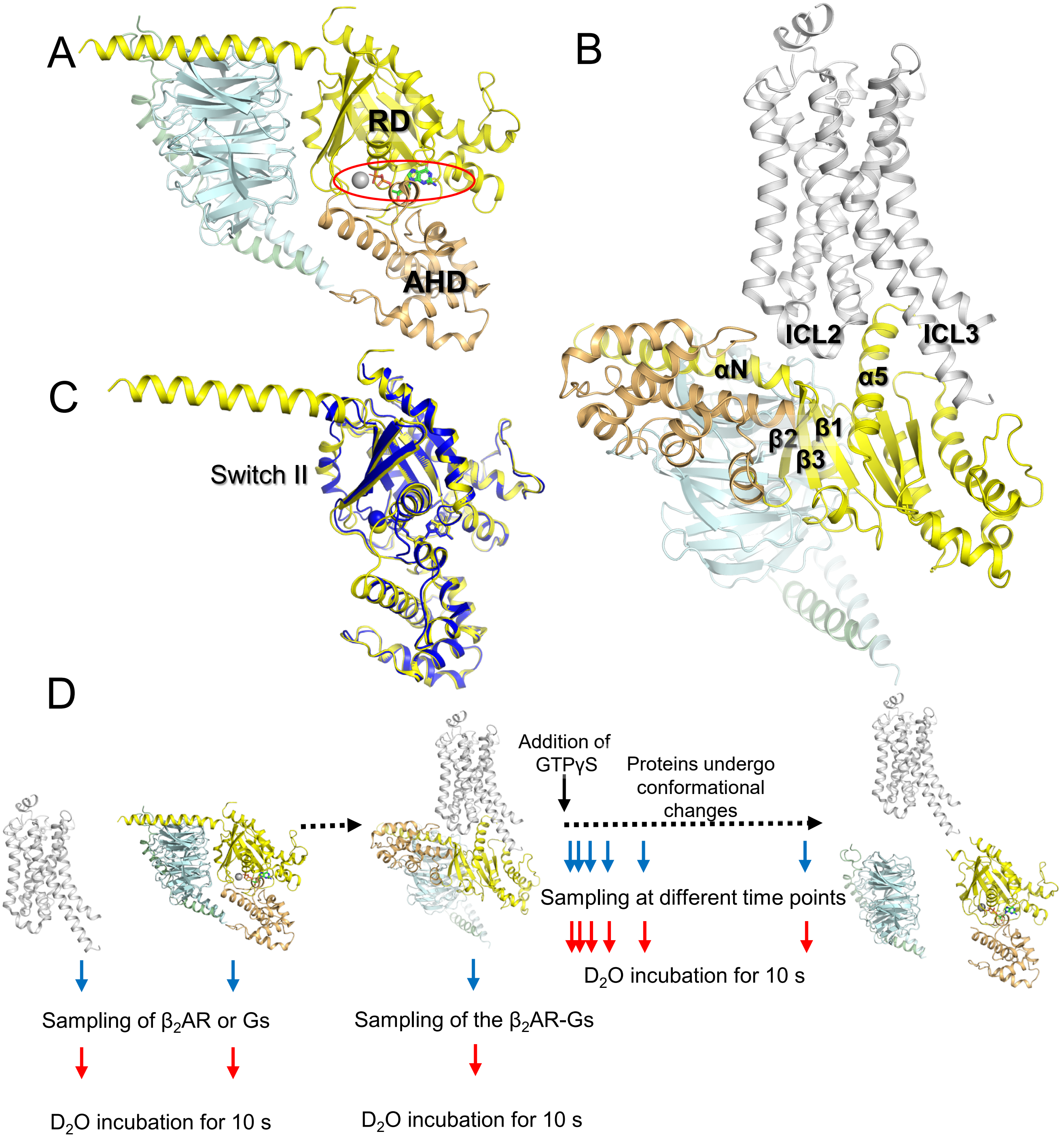


**Figure S1. Structures representing different stages of the G protein activation cycle and scheme of the pulse-labeling HDX-MS; Related to Figures 1 and 2.**

(A) X-ray crystal structure of the GDP-bound heterotrimeric Gs protein (PDB: 6EG8). GDP is shown as sticks, and $Mg^{2+}$ is indicated with a gray sphere. The Ras-like GTPase domain (RD) of Gα is colored in yellow, the α-helical domain (AHD) of Gα is in light orange, Gβ is in light blue, and Gγ is in light green. The GDP- or GTP-binding region (i.e., the nucleotide-binding pocket) is located between the RD and AHD (red circle). (B) X-ray crystal structure of the nucleotide-free $\beta_2$AR–Gs complex (PDB: 3SN6). $\beta_2$AR is shown in gray. The RD of Gα is colored in yellow, AHD of Gα is in light orange, Gβ is in light blue, and Gγ is in light green. Nanobody Nb35 and the T4 lysozyme insertion are omitted for clarity. The $\beta_2$AR–Gs complex shows the receptor–Gs interfaces and a large movement of AHD. (C) Comparison of Gαs in the GDP-bound heterotrimeric state (yellow, PDB: 6EG8) and GTPγS-bound state (blue, PDB: 1AZT). Gβ and Gγ are omitted for clarity. GDP and GTPγS are shown as sticks, and $Mg^{2+}$ is indicated with a sphere. (D) Schematic of the pulse-labeling HDX-MS protocol. To analyze the time-resolved conformational changes of the nucleotide-free $\beta_2$AR–Gs complex, the protein is triggered (for example, by addition of GTPγS to the nucleotide-free $\beta_2$AR–Gs complex) to undergo conformational changes, followed by a $D_2O$ pulse for a short period (10 s) at specific time points.

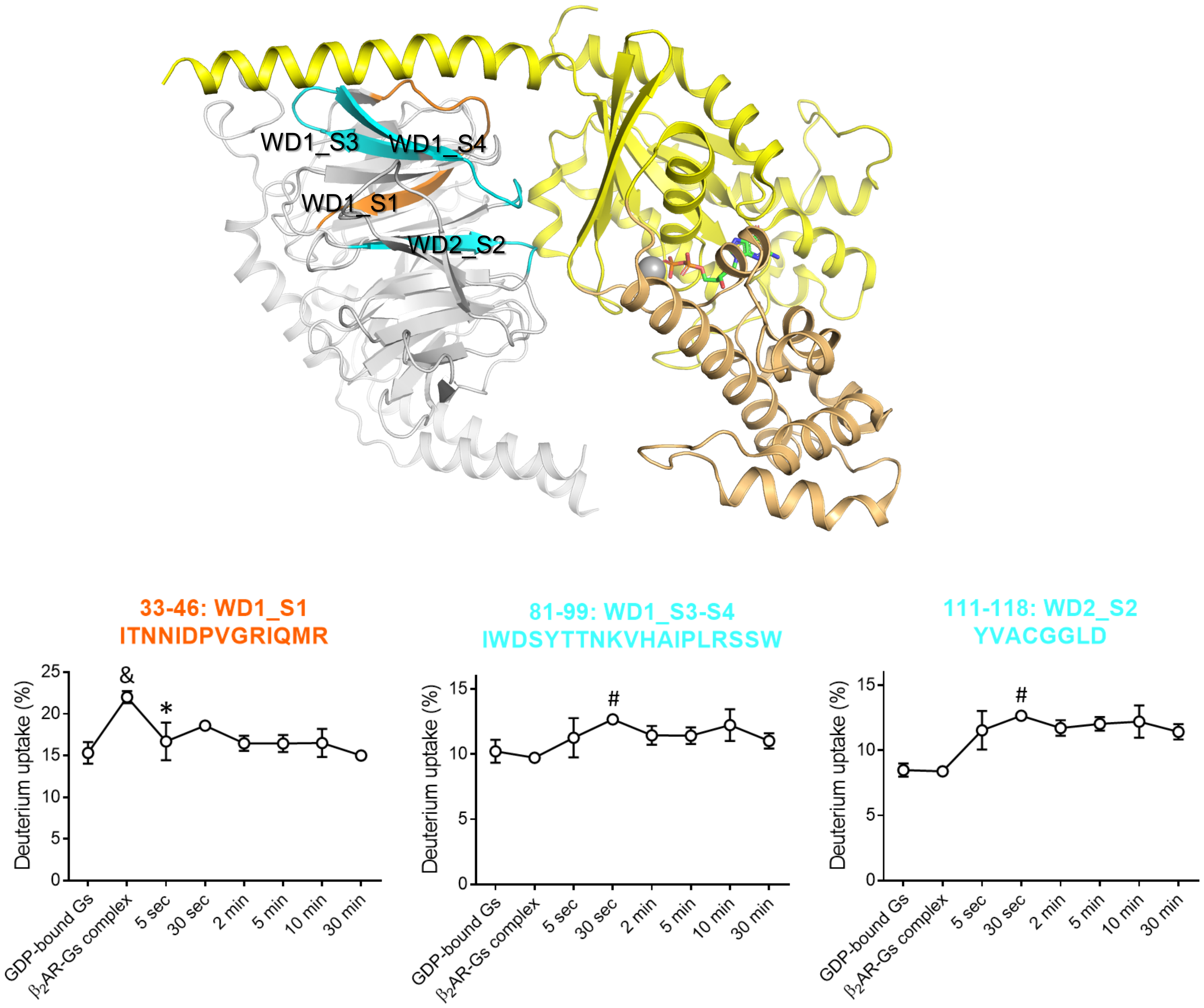


**Figure S2. Time-resolved analysis of Gβ after GTPγS addition to the $β_2$AR–Gs complex; Related to Figure 1.**

Upper panel: Regions in Gβ showing HDX profile changes after GTPγS addition are color-coded on the X-ray crystal structure of the GDP-bound Gs heterotrimer (PDB: 6EG8). Gαs is shown in yellow (RD) and orange (AHD). Lower panel: Pulse-labeling deuterium uptake plots for the color-coded peptic peptides. A mass difference >0.3 Da was considered significant. To compare two different time points, a paired *t*-test was used, and $p < 0.05$ was considered to be statistically significant. &, the HDX level of the nucleotide-free $β_2$AR-Gs complex is significantly different from that of GDP-bound Gs. *, the first time point after GTPγS addition when the HDX level returned to that of the GDP-bound Gs. #, the first time point after GTPγS addition that showed a significant difference from the $β_2$AR–Gs complex. Error bars represent the standard error of the mean of more than three independent experiments. Data are plotted using a non-linear/non-logarithmic scale.

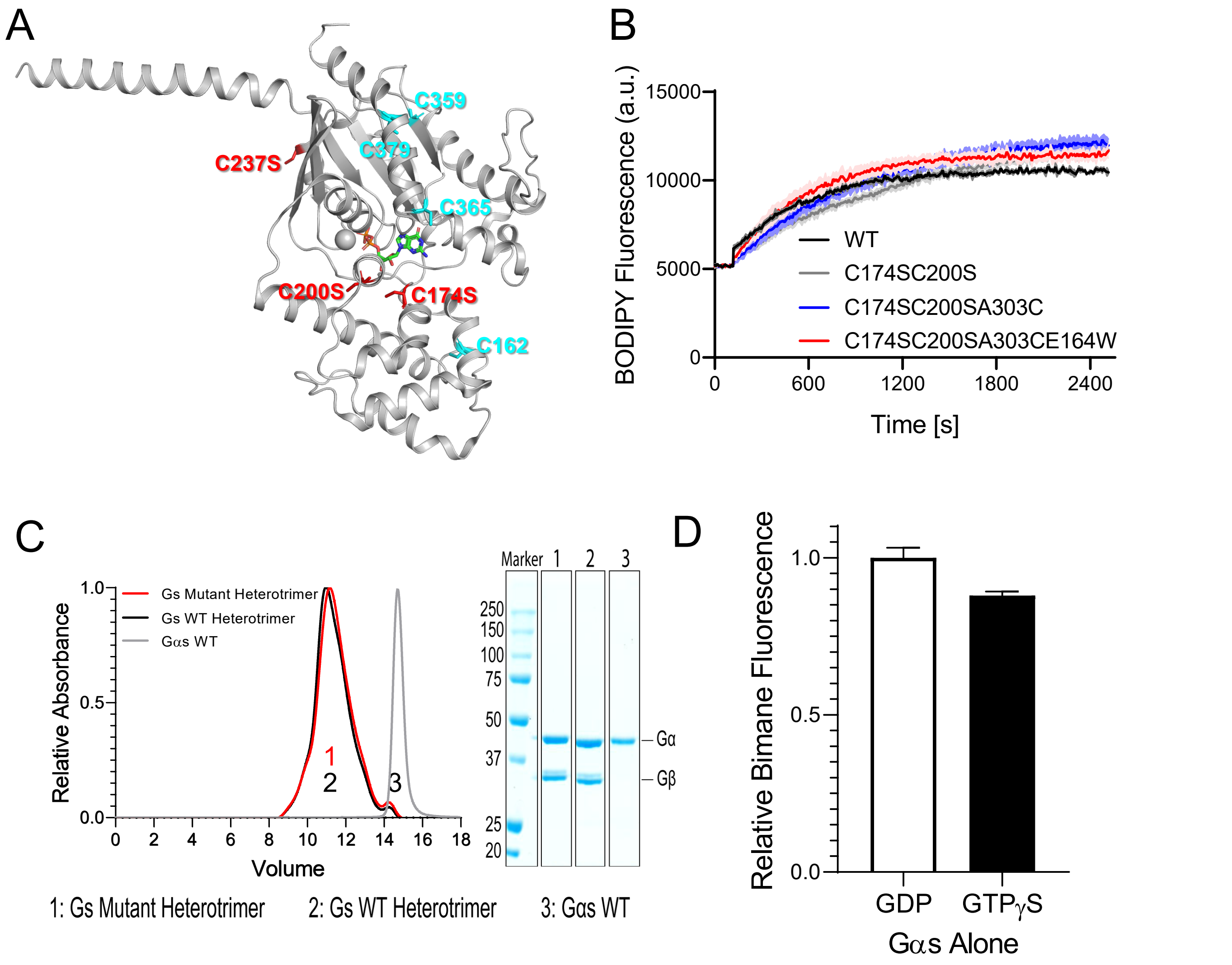


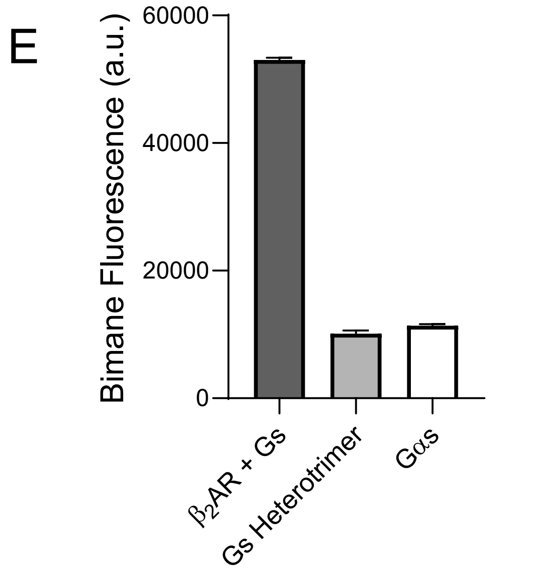


**Figure S3. Development of a tryptophan-induced bimane quenching system to monitor the movement of the AHD; Related to Figures 2 and 4.**
(A) The cysteine residues in Gαs either exposed to the solvent (red) or excluded from the solvent (cyan) in GDP-bound Gαs are indicated as sticks on the X-ray crystal structure of the GDP-bound heterotrimeric Gs protein (PDB: 6EG8). Gβγ is omitted for clarity. (B and C) Functional intactness of the mutant constructs tested by BODIPY-FL-GTPγS incorporation (B) and heterotrimer formation with Gβγ by analyzing size-exclusion chromatography (C). (D) Relative bimane fluorescence in GDP- or GTPγS-bound Gαs. (E) Bimane fluorescence in the nucleotide-free $\beta_2$AR–Gs complex is higher than that in the GDP-bound Gs heterotrimer or GDP-bound Gαs, indicating that the bimane fluorescence is quenched in the GDP-bound state but not in the nucleotide-free $\beta_2$AR–Gs complex due to a larger distance between the RD and AHD in the nucleotide-free $\beta_2$AR–Gs complex than in the GDP-bound state. Error bars represent the standard error of the mean of three independent experiments.

A

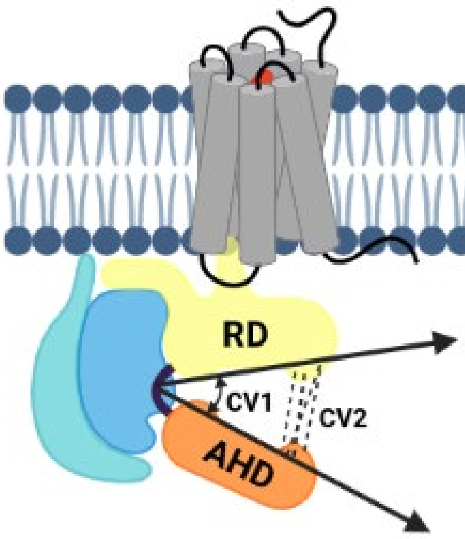


B

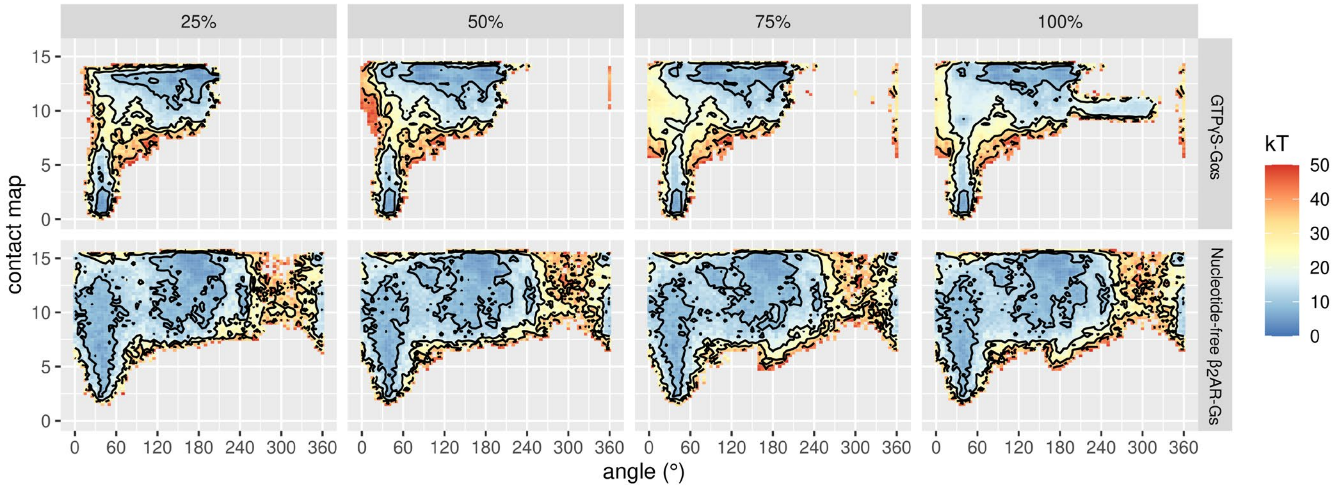


**Figure S4. Enhanced conformational sampling of AHD movement by metadynamics simulations; Related to Figure 3.**

(A) Collective variables (CV1 and CV2) used for metadynamics simulations of GTPγS-bound Gαs and the nucleotide-free $\beta_2$AR–Gs complex. The cartoon was generated with BioRender. (B) Reconstructed free energy of Gαs rearrangement between closed and open conformations in the GTPγS–Gαs and nucleotide-free $\beta_2$AR–Gs systems from metadynamics simulations as a function of the angle between the RD and AHD (horizontal axis; see definition in the STAR Methods) and deviation of the RD–AHD contact map based on PDB 1AZT (vertical axis; see definition in STAR Methods). The color bar is in units of kT. Data are plotted for the first-quarter, half, three-quarters, and whole trajectories in columns and for the two systems studied in rows.

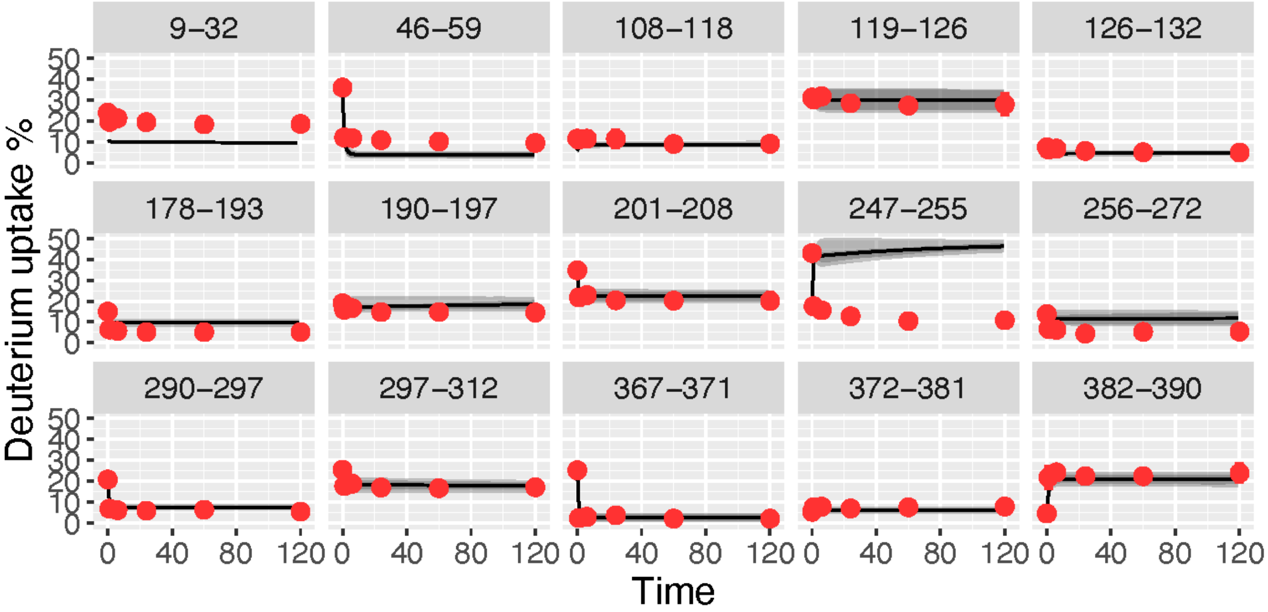


**Figure S5. Comparison between predicted and measured values of deuterium uptake; Related to Figure 3.**

Comparison between the experimental deuterium uptake values measured at different times for different peptide segments from Gαs (see Figures 1 and 2 for corresponding amino acid sequences and locations) and the predicted values $D_J(t)$ from the posterior distribution of the kinetic model built as described in the STAR Methods section (eq. 7). The red dots refer to experimental values and the black lines correspond to the averages of the sampled values according to the kinetic model. Gray areas demarcate the credible interval corresponding to the 25% and 75% quantiles.

A

| Prey ID | Description | Reporter expression | | |
|---|---|---|---|---|
| | | *lacZ* | *URA3* | *ADE2* |
| **AD-Hybrid - 1** | The activation domain (AD) is fused in frame to the 260th aa of MAGE family member D2 (**MAGE D2**) (NM_014599). | + | + | + |

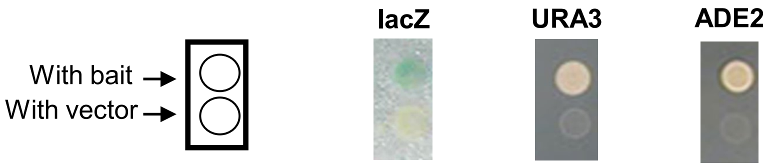


B

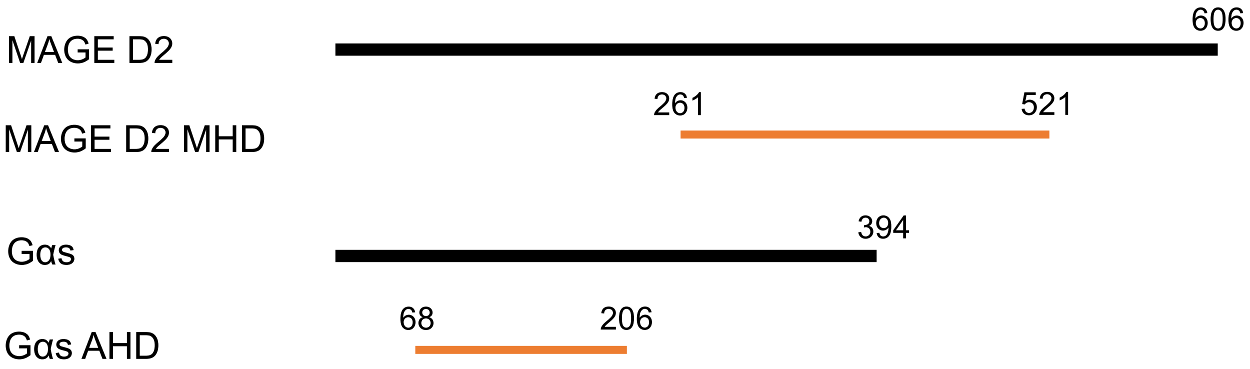


**Figure S6. Identification of MAGE D2 as a novel Gαs AHD-binding protein; Related to Figure 4.**

(A) Yeast-two-hybrid library screening of a human kidney cDNA AD library using the Gαs AHD as bait identified MAGE D2 (NM_014599) as a Gαs AHD-binding protein (see STAR Methods for details). (B) The protein constructs of MAGE D2 MHD and Gαs AHD generated for experiments in Figure 4C and 4D (see STAR Methods for details).

```
                                                     αN
sp|P63092|GNAS2_HUMAN      --------MGCLGNSK---------TEDQRNEEKAQREANKKIEKQLQKDKQVYRATHRL    43
sp|Q03113|GNA12_HUMAN      MSGVVRTLSRCLLPAEAGGARERRAGSGARDAERAAARRRSRDIDALLARERRAVRRLVKI    60
sp|P63096|GNAI1_HUMAN      --------MGCTLS----------------AEDKAAVERSKMIDRNLREDGEKAAREVKL    36
sp|P50148|GNAQ_HUMAN       --MTLESIMACCLS----------------EEAKEARRINDEIERQLRRDKRDARRELKL    42
                                     *                      :   . .  *:  * .: .      ::

                           β1            α1                                     αA
sp|P63092|GNAS2_HUMAN      LLLGAGESGKSTIVKQMRILHVNGFNGEGGEEDPQAARSNSDGEKATKVQDIKNNLKEAI   103
sp|Q03113|GNA12_HUMAN      LLLGAGESGKSTFLKQMRIIHGREF---------------DQKALLEFRDTIFDNILKGS   105
sp|P63096|GNAI1_HUMAN      LLLGAGESGKSTIVKQMKIIHEAGY---------------SEEECKQYKAVVYSNTIQSI    81
sp|P50148|GNAQ_HUMAN       LLLGTGESGKSTFIKQMRIIHGSGY---------------SDEDKRGFTKLVYQNIFTAM    87
                           ****:*******::***:*:*   :               .  :         : .*  .

                                                       αB                    αC
sp|P63092|GNAS2_HUMAN      ETIVAAMSNLVPPVELANPENQFRVDYILSVM---NVPDF-DFPPEFYEHAKALWEDEGV   159
sp|Q03113|GNA12_HUMAN      RVLVDARDKLGIPWQYSENEKHGM--FLMAFENKAGLPVEPATFQLYVPALSALWRDSGI   163
sp|P63096|GNAI1_HUMAN      IAIIRAMGRLKIDFGDSARADDAR--QLFVLA---GAAEEGFMTAELAGVIKRLWKDSGV   136
sp|P50148|GNAQ_HUMAN       QAMIRAMDTLKIPYKYEHNKAHAQ--LV-REV---DVEKVSAFENPYVDAIKSLWNDPGI   141
                            .:: * . *           .     :       .               . **.* *:

                                αD              αE             αF              β2
sp|P63092|GNAS2_HUMAN      RACYERSNEYQLIDCAQYFLDKIDVIKQADYVPSDQDLLRCRVLTSGIFETKFQVDKVNF   219
sp|Q03113|GNA12_HUMAN      REAFSRRSEFQLGESVKYFLDNLDRIGQLNYFPSKQDILLARKATKGIVEHDFVIKKIPF   223
sp|P63096|GNAI1_HUMAN      QACFNRSREYQLNDSAAYYLNDLDRIAQPNYIPTQQDVLRTRVKTTGIVETHFTFKDLHF   196
sp|P50148|GNAQ_HUMAN       QECYDRRREYQLSDSTKYYLNDLDRVADPAYLPTQQDVLRVRVPTTGIIEYPFDLQSVIF   201
                           : .:.*  *:** :.. *:*:.:* : :  *.*:.**:*  *  *.**.*  * ...: *

                              β3           α2          β4                       α3
sp|P63092|GNAS2_HUMAN      HMFDVGGQRDERRKWIQCFNDVTAIIFVVASSSYNMVIREDNQTNRLQEALNLFKSIWNN   279
sp|Q03113|GNA12_HUMAN      KMVDVGGQRSQRQKWFQCFDGITSILFMVSSSEYDQVLMEDRRTNRLVESMNIFETIVNN   283
sp|P63096|GNAI1_HUMAN      KMFDVGGQRSERKKWIHCFEGVTAIIFCVALSDYDLVLAEDEEMNRMHESMKLFDSICNN   256
sp|P50148|GNAQ_HUMAN       RMVDVGGQRSERRKWIHCFENVTSIMFLVALSEYDQVLVESDNENRMEESKALFRTIITY   261
                           :*.******.:*:**::**:.:*:*:* *: *.*: *: *. . **: *:  :* :* .

                                     β5          αG                                α4
sp|P63092|GNAS2_HUMAN      RWLRTISVILFLNKQDLLAEKVLAGKSKIEDYFPEFARYTTPEDATPEPGEDPRVTRAKY   339
sp|Q03113|GNA12_HUMAN      KLFFNVSIILFLNKMDLLVEKVKT--VSIKKHFPDFRGDP------------HRLEDVQR   329
sp|P63096|GNAI1_HUMAN      KWFTDTSIILFLNKKDLFEEKIKK--SPLTICYPEYAGSN------------TYEEAA-A   301
sp|P50148|GNAQ_HUMAN       PWFQNSSVILFLNKKDLLEEKIMY--SHLVDYFPEYDGPQ------------RDAQAARE   307
                             :   *:****** **: **:      :   :*::                   :  .

                                                 β6                  α5
sp|P63092|GNAS2_HUMAN      FIRDEFLRISTASGDGRHYCYPHFTCAVDTENIRRVFNDCRDIIQRMHLRQYELL        394
sp|Q03113|GNA12_HUMAN      YLVQCFD-RKR--RNRSKPLFHHFTTAIDTENVRFVFHAVKDTILQENLKDIMLQ        381
sp|P63096|GNAI1_HUMAN      YIQCQFEDLNK--RKDTKEIYTHFTCATDTKNVQFVFDAVTDVIIKNNLKDCGLF        354
sp|P50148|GNAQ_HUMAN       FILKMFVDLNP--D-SDKIIYSHFTCATDTENIRFVFAAVKDTILQLNLKEYNLV        359
                           ::   *   .       :  : *** * **:*:: **    * * : :*::  *
```

**Figure S7. Sequence alignment of Gαs, Gαi1, Gαq, and Gα12; Related to Figure 4.**
The sequence similarities between Gαs, Gαi1, Gαq, and Gα12 are analyzed by the multiple sequence alignment Clustal Omega program (http://www.clustal.org/omega/). The linkers between the RD (black) and AHD (orange) regions are colored in blue. The "*" symbol indicates single, fully conserved residues; the ":" symbol indicates conservation between groups of strongly similar properties; and the "." symbol indicates conservation between groups of weakly similar properties.

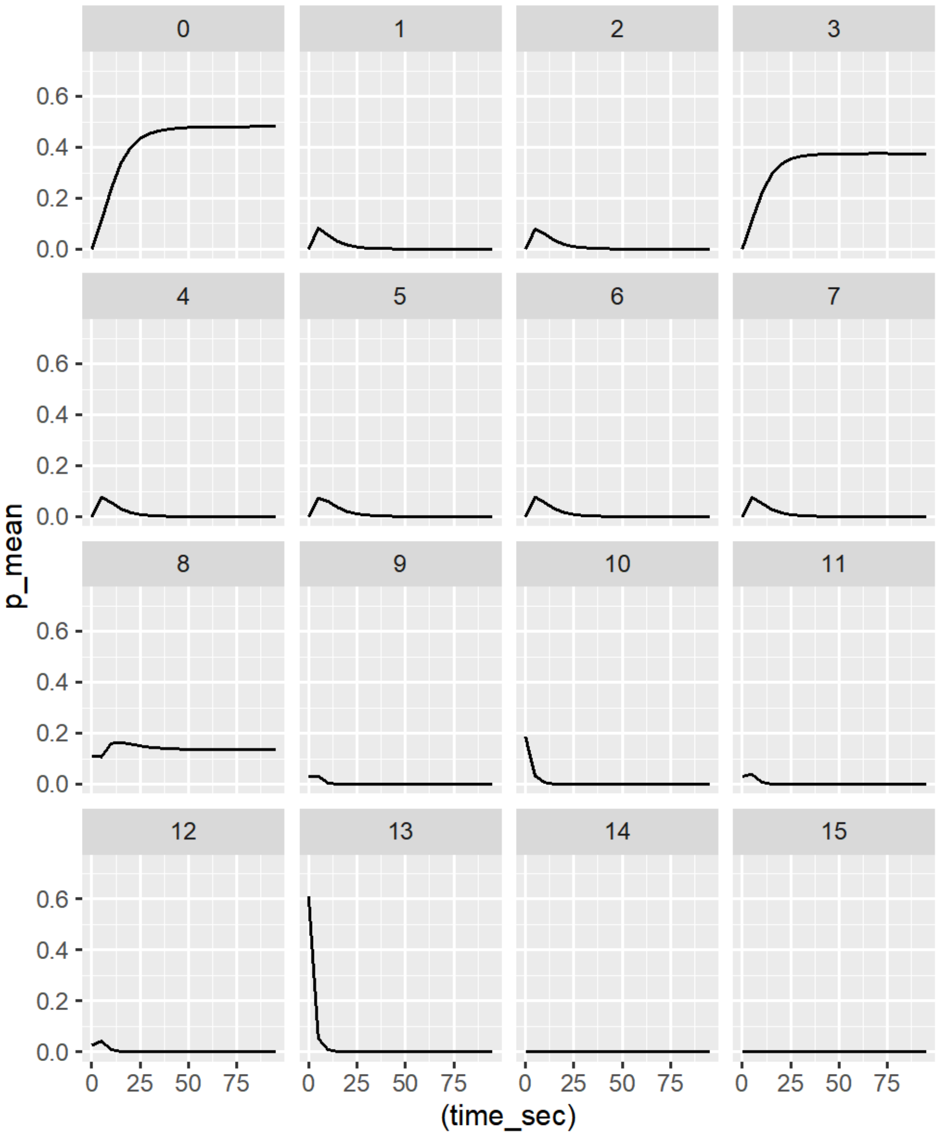


**Figure S8. Time evolution of the probability of each identified conformational cluster of Gαs from simulations; Related to Figure 3.**

Time evolution of the probability of each identified conformational cluster according to the posterior distribution of the transition matrix ($\boldsymbol{\pi}(t)$ in eq. 9). The black line corresponds to the average of the sampled values. The credible intervals corresponding to the 25% and 75% quantiles are reported in gray, but are indistinguishable from the mean values on the scale of the plot.

**Table S1.** Initial and equilibrium probabilities of highly populated conformational states (maximum probability larger than 10% across time) according to the proposed kinetic model. Credible intervals are obtained from the 25% and 75% quantiles of the posterior distribution.

| State | Initial probability | Initial probability C.I. | Equilibrium probability | Equilibrium probability C.I. |
|---|---|---|---|---|
| 0 | 0% | - | 29% | (9%,48%) |
| 3 | 0% | - | 51% | (37%,62%) |
| 5 | 0% | - | 13% | (0%,28%) |
| 10 | 29% | (18%,40%) | 0% | - |
| 13 | 54% | (45%,63%) | 0% | - |

**Table S2.** Mean-first passage times between highly probable states (maximum probability larger than 10% across time) according to the proposed kinetic model. MFPT are reported in seconds, along with 25% credible intervals from the posterior sampling. Only transitions with MFPT shorter than 1000 sec are reported.

| End state | Start State | MFPT (sec) | (25%-quantile, 75% quantile) |
|---|---|---|---|
| 0 | 3 | 141.9 | (0.2,273.5) |
| 0 | 5 | 140.3 | (0.4,271.5) |
| 0 | 10 | 134.0 | (0.5,261.0) |
| 0 | 13 | 134.4 | (0.4,261.0) |
| 3 | 0 | 299.1 | (0.3,579.0) |
| 3 | 5 | 21.6 | (0.4,49.6) |
| 3 | 10 | 27.4 | (0.5,54.2) |
| 3 | 13 | 26.9 | (0.5,54.2) |
| 5 | 3 | 224.6 | (82.5,250.7) |
| 5 | 10 | 167.3 | (60.2,166.9) |
| 5 | 13 | 164.4 | (58.2,164.5) |

**Table S3.** Average RMSD values of the HDX-MS peptides between macrostates 3 and 13, and 3 and 10, respectively.

| Fragment | Region | 3 vs 13 | 3 vs 10 |
|---|---|---|---|
| 46-59 | p-loop – α1 | 2.60 | 1.42 |
| 108-118 | N term of αA/αB loop | 1.61 | 0.57 |
| 119-126 | αA/αB loop - αB | 1.12 | 0.85 |
| 126-132 | αB | 0.31 | 0.39 |
| 178-193 | αE | 0.66 | 0.49 |
| 190-197 | αE/αF loop | 0.57 | 0.32 |
| 201-208 | Switch I | 2.56 | 2.24 |
| 256-272 | Switch III | 2.04 | 1.58 |
| 290-297 | β5 – αG | 0.42 | 0.44 |
| 297-312 | αG – αG/α4 | 1.71 | 1.61 |
| 367-371 | β6/α5 – N-terminus of α5 | 1.95 | 1.84 |
| 372-381 | N-terminal half of a5 | 0.30 | 0.33 |
| 382-390 | C-terminal half of a5 | 0.39 | 0.40 |
